# Integrated analysis sheds light on evolutionary trajectories of young transcription start sites in the human genome

**DOI:** 10.1101/192757

**Authors:** Cai Li, Boris Lenhard, Nicholas M. Luscombe

## Abstract

Previous studies revealed widespread transcription initiation and fast turnover of transcription start sites (TSSs) in mammalian genomes. Yet how new TSSs originate and how they evolve over time remain poorly understood. To address these questions, we analyzed ∼200,000 human TSSs by integrating evolutionary and functional genomic data, particularly focusing on TSSs that emerged in the primate lineages. We found that intrinsic factors of repetitive sequences and their proximity to established regulatory modules (extrinsic factors) contribute significantly to origin of new TSSs. In early periods, young TSSs experience rapid sequence evolution driven by endogenous mutational mechanisms that reduce the instability of associated repetitive sequences. In later periods, the regulatory functions of young TSSs are gradually modified, and with evolutionary changes subject to temporal (fewer regulatory changes in younger TSSs) and spatial constraints (fewer regulatory changes in more isolated TSSs). These findings advance our understanding of how regulatory innovations arise in the genome throughout evolution and highlight the roles of repetitive sequences in these processes.

## 1. Introduction

Many studies revealed that transcription is pervasive in prokaryotic and eukaryotic genomes^1,2^. One recent study found that three-quarters of the human genome can be transcribed^3^, indicating a much more complex transcriptional landscape than previously thought. Transcription Start Sites (TSSs) are the genomic loci where transcription initiation occurs and thus are a critical class of regulatory element for transcriptional control. By harnessing diverse high-throughput sequencing technologies, studies in the past few years have greatly improved TSS annotation in model organism genomes, especially human, and uncovered new characteristics of transcriptional initiation^4-6^. One intriguing phenomenon about TSSs is that they occur widely throughout the genome, not only in typical promoters of annotated genes, but also in other regions such as intergenic or intronic loci. For example, some enhancers also contain TSSs, producing so-called enhancer RNAs^7-9^.

Many previous studies about TSS evolution focused on cross-species comparisons and revealed interesting macro-evolutionary patterns^10-14^. For example, by comparing human and mouse TSSs, a recent study found that >56% of protein-coding genes have experienced TSS turnover events since humans and mice diverged^13^. Genes with TSS turnover were also found to experience adaptive evolution in their coding regions and expression levels^13^. Unlike macro-evolution, however, micro-evolutionary processes (i.e. intra-species evolution) of TSSs are relatively poorly understood. Given the high turnover rate of TSSs^13^, population genomic data can provide a more detailed view of TSS evolution. Although some previous studies made use of population genomic data, they pooled all TSSs together to compare with non-TSS elements^15^ or focused on purifying selection^13,16^. Since different TSSs could have distinct evolutionary histories, pooling all TSSs together could bury the interesting characteristics of a specific TSS categories. A recent comprehensive study in *Drosophila melanogaster* populations investigating the relationship between genetic variations and TSS usage identified thousands of genetic variants affecting transcript levels and promoter shapes, providing important new insights into TSS evolution at the population level^17^.

Despite extensive investigation, many questions about TSSs are yet to be addressed. Importantly, the evolutionary origin of new TSSs and evolutionary trajectories of newly emerged TSSs remain unresolved. Previous studies have suggested that repetitive sequences are a rich source of new TSSs^13,18^, but the underlying mechanisms of how these sequences contribute to novel transcription initiation remain unclear. For instance, why do some repetitive elements initiate transcription and others not? How does the host genome handle the potential conflicts arising from the inherent instability of repetitive elements associated with new TSSs? Furthermore, the subsequent changes of newly emerged TSSs and their evolutionary fates have not been systematically investigated. Only by addressing these questions can we begin to understand how regulatory innovations arise in the genome throughout evolution and how they contribute to biological diversity and adaptation.

To gain detailed insights into evolution of young TSSs and the underlying regulatory mechanisms, we analyzed ∼200,000 published human TSSs by integrating both evolutionary (inter-species and intra-species) and functional genomic approaches, with an emphasis on evolutionarily young TSSs that emerged in the primate lineages. We show that 1) intrinsic factors of repetitive sequences and extrinsic chromatin environments contribute significantly to the origin of novel transcription initiation; 2) after emerging in the genome, young TSSs undergo rapid sequence evolution which is likely due to several endogenous mutational mechanisms; and 3) regulatory outcomes of young TSSs are gradually modified in subsequent periods and tend to be subject to temporal and spatial constraints.

## 2. Results

### 2.1 Identification of evolutionarily young TSSs in the human genome

Using the cap analysis of gene expression (CAGE) sequencing technologies, the FANTOM 5 project^4^ generated the most comprehensive TSS annotation to date, covering major primary cell types and tissues in human. To identify evolutionarily young TSSs, we took advantage of the ‘robust’ human TSS dataset from FANTOM project, which consists of 201,873 high-confidence TSSs. After filtering TSSs that could confound downstream analysis (see Methods for details), we grouped the remaining 151,902 TSSs into categories of different evolutionary ages. Since there is no large-scale CAGE TSS annotation in the other primate genomes, it is impossible to define the evolutionary ages of TSSs by comparing TSS annotations. However, previous studies revealed that sequence-intrinsic properties of many promoters can drive transcription initiation autonomously^19,20^, indicating that the sequence itself is an important determinant of promoter capacity. Moreover, Young et al. (2015) found that, of those human TSSs that could be aligned to an orthologous sequence in the mouse, more than 80% have detectable transcriptional initiation in mouse^13^. This implies that if the orthologous sequence of a human TSS can be found in another genome, it probably exhibits initiation in that species.

Therefore, to estimate the evolutionary ages of human TSS loci, we investigated the sequence presence/absence patterns based on sequence alignments between human and other 16 genomes (10 primate species representing major primate lineages and 6 non-primate mammalian species as outgroups). A human TSS locus is considered present in another genome if the corresponding pairwise alignment satisfies: 1) a mapping ratio of the human TSS peak (i.e. a CAGE tag cluster region predicted by decomposition-based peak identification method in FANTOM) in another genome of ≥90% and 2) a mapping ratio of the TSS peak±100 bp (considered as core promoter region in this study) of ≥50% (see Methods and **Supplementary Tables 1-3** for more details). Based upon the presence/absence patterns in alignments, we categorized the human TSSs into four groups of different sequence ages (**Fig. 1a**): 1) TSSs whose sequence loci can be found in at least one non-primate mammalian genome, consisting of 141,117 TSSs (92.9% of all surveyed TSSs, named ‘mammalian’ group; **Fig. 1a**); 2) TSSs whose sequences occurred during early primate evolution but before the last common ancestor of Old World anthropoids, consisting of 6,668 TSSs (4.4%, named ‘primate’ group; **Fig. 1a**); 3) TSSs whose sequences occurred during the evolution of Old World anthropoids but before the last common ancestor of hominids, consisting of 3,318 TSSs (2.2%, named ‘OWA’ group; **Fig. 1a**); 4) TSSs whose sequences occurred since emergence of hominids, consisting of 799 TSSs (0.5%, named ‘hominid’ group; **Fig. 1a**). The relatively large numbers of TSSs in three recent periods corroborate the “frequent birth” phenomenon reported previously^13^, and enable us to perform detailed comparative analysis between these periods. Hereafter we considered TSSs in the ‘mammalian’ group as evolutionarily old TSSs and those in other three groups as evolutionarily young TSSs. For instance, in the gene *BAAT* locus shown in **Fig. 1b**, there are two old TSSs present in both primate and non-primate mammalian genomes, and one young TSS established during the evolution of OWAs. The young TSS is located in a region overlapping one long terminal repeat (LTR) element (**Fig. 1b**), suggesting that it originated from an LTR insertion. This young TSS is expressed in many cell types where the old TSSs are expressed, suggesting it may undertake part of the transcription task of old TSSs or up-regulate the expression level of *BAAT* in some conditions.

**Fig. 1.**
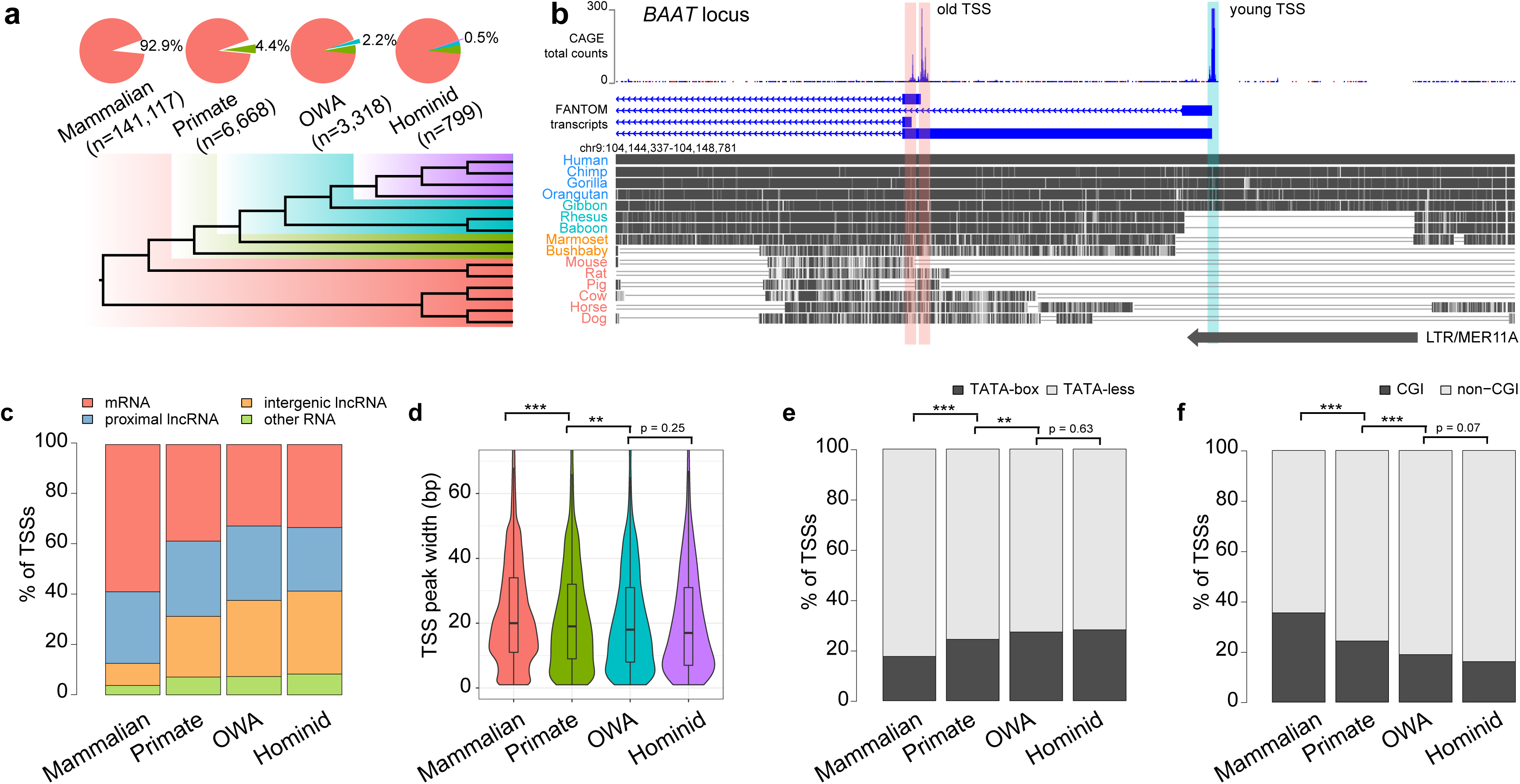
Classification of human TSSs by evolutionary age. (**a**) Statistics of four TSS groups defined by sequence age using genomic alignments. At the bottom is the phylogeny with colors indicating the corresponding period of each TSS group. (**b**) An example gene locus shows two ‘mammalian’ TSSs (red shade) and one ‘OWA’ TSS (cyan shade). An LTR element overlapping the young TSS can be seen at the bottom of the alignment. CAGE tag counts and transcript isoforms shown at the top were from FANTOM CAT annotation (part of FANTOM 5). Genome alignments represented by grey blocks and lines were generated using UCSC genome browser (hg19). (**c**) Composition of transcription type in each TSS group. Transcript types are derived from FANTOM CAT annotation. (**d**) Violin and box plots for TSS peak widths of each TSS group. (**e**) Proportions of TATA-box containing and TATA-less TSSs. (**f**) Proportions of CGI-associated and non-CGI-associated TSSs. Statistical significances in panel **d** were calculated by one-tailed Wilcoxon rank sum tests; statistical significances in panels **e** and **f** by Fisher’s exact tests; “**”, p < 0.01; “***”, p < 0.001.

We first examined some general features among TSS groups. We found that old TSSs are mainly associated with mRNAs (59%), while many young TSSs are associated with lncRNAs (54%∼60%), indicating a compositional bias in the TSS groups (**Fig. 1c**). As TSSs become older, the proportion of mRNA TSSs becomes larger, and the opposite happens to the intergenic lncRNA TSSs (**Fig. 1c**). Relative to older TSSs, younger TSSs generally have narrower TSS peaks (**Fig. 1d**) and comprise more TATA-box containing TSSs (**Fig. 1e**) and fewer CpG island (CGI)-associated TSSs (**Fig. 1f**). This is consistent with previous observations about broad and sharp promoters in mammalian genomes^4,21^, which found that CGI promoters are usually broad and associated with housekeeping genes, while TATA-box promoters are sharp and associated with less conserved tissue-specific genes. Both old and young TSSs exhibit elevated GC content and CpG content in TSS-proximal positions (**Supplementary Fig. 1**), although relative to young TSSs, old TSSs tend to be more GC-rich. We also noticed that the ‘hominid’ TSS group has higher average GC and CpG content relative to ‘OWA’ and ‘primate’ groups (**Supplementary Fig. 1**), which could be partly due to fewer historical deamination events of methylated cytosines in very young TSS loci (see also later sections about DNA methylation).

### 2.2 Sources of young TSSs

#### 2.2.1 Intrinsic factors of repetitive sequences contribute to novel transcription initiation

Based upon the defined TSSs groups of different ages, next we systematically investigated how new TSSs originate and how they evolve over time. Previous analyses from earlier FANTOM projects showed that many mammalian transcripts initiate within repetitive elements, especially retrotransposons^13,18^. Given the extensive retrotransposition during mammalian evolution, retrotransposon-derived TSSs could be an important source of novel TSSs. In addition, tandem repeats, which are highly mutable loci, were found to be abundant in promoter regions and have significant impact on gene expression^22,23^. With these observations in mind, we examined the repetitive sequences (or ‘repeats’ for short hereafter) in all TSS loci, including transposable elements (TEs, i.e. retrotransposons and DNA transposons) and tandem repeats, based on annotations of RepeatMasker^24^, TRF^25^ and STRcat^26^. We found that ∼70% of young TSSs have at least one repeat element within core promoter regions (±100 bp of TSSs), but only 24% among old TSSs (**Fig. 2a**). Whereas a large fraction (43%) of repetitive sequences associated with old TSSs are tandem repeats, many young TSS loci are associated with retrotransposons, including LTRs, long intersperse nuclear elements (LINEs) and short interspersed nuclear elements (SINEs) (**Fig. 2a**). Because some tandem repeats could derive from retrotransposons, we performed an alternative analysis considering only the nearest retrotransposon element (**Supplementary Table 4 & Supplementary Fig. 2**). LTRs are the most abundant retrotransposon class associated with young TSSs, with ∼30% of young TSSs are associated with LTRs. 14% and 8% of young TSSs are associated with LINEs and SINEs, respectively. The large number of retrotransposons associated with young TSSs suggests a major role of retrotransposition in forming new TSS loci.

**Fig. 2.**
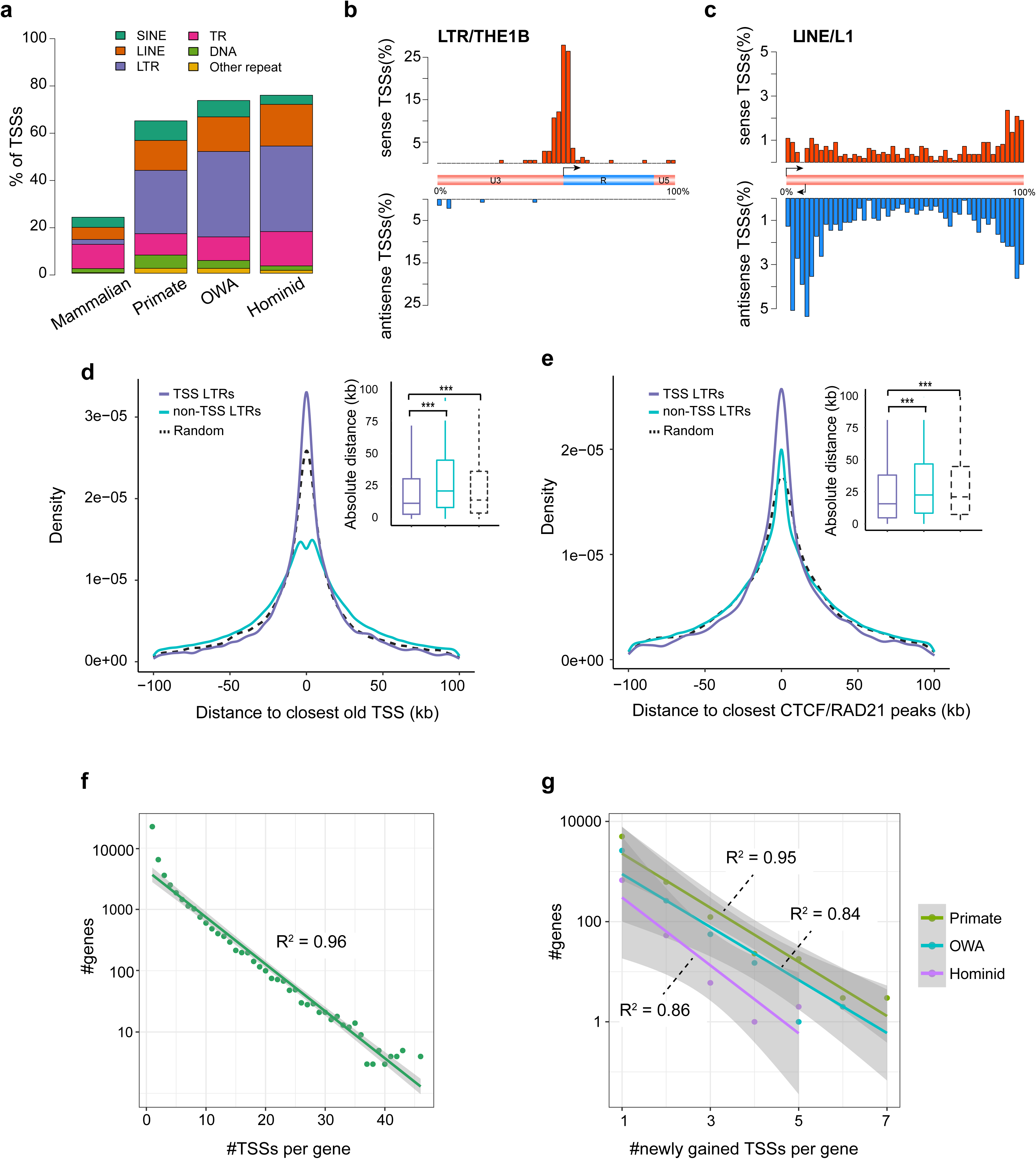
Intrinsic and extrinsic factors contributing to the origin of new TSSs. (**a**) Composition of major repeat families in four TSS groups. To obtain a non-redundant assignment, we considered the nearest repeat element within TSS±100 bp. (**b**) Distribution of young TSSs along the LTR/THE1B elements, with a bin size of 2% of its full-length consensus sequence. In the middle is the THE1B structure, which includes the original TSS, U3, R and U5 regions for the transposable element. (**c**) Distribution of young TSSs along the LINE/L1 elements, with a bin size of 2% of full-length consensus sequences. In the middle is the L1 structure, which indicates the sense and antisense L1 TSSs at 5’end. (**d**) Comparison of distances of TSS-associated and non-TSS-associated LTRs to the closest old TSSs. The distances of random intervals (generated by “bedtools shuffle” with TSS-associated LTRs as input) to the closest old TSSs are also provided for comparison. (**e**) Comparison of distances of TSS-associated and non-TSS-associated LTRs to the closest CTCF or RAD21 ChIA-PET peaks (GM12878). Random intervals used here is the same as that in panel **d**. (**f**) Exponential approximation for the number of genes with a certain number of TSSs and number of TSSs per gene, based on data of all TSSs. R^2^ is the coefficient of determination for the linear regression in the figure. (**g**) Exponential approximation for the number of genes and number of newly gained TSSs per gene, based on data of newly emerged TSSs in three periods. R^2^ is the coefficient of determination. Statistical significances in panels **d** and **e** were calculated by one-tailed Wilcoxon rank sum tests; “***”, p < 0.001.

Faulkner et al. (2009) revealed that many TE-derived TSSs are unevenly distributed along TE element consensus sequences, and many TE-derived TSSs are not present in the canonical 5’ promoters of TE elements^18^. However, how these TE-derived sequences contribute to transcription initiation was not discussed in detail and thus remain poorly understood. To gain more detailed insight into this question, we first mapped TSSs to the TE consensus sequences like Faulkner et al. (2009), and analyzed the distributions of TSSs along repeat elements. The distributions obtained from our analysis are similar to those in Faulkner et al. (2009), but also exhibit some differences. The differences are likely due to the upgraded CAGE protocols^27^ and improvements in the TSS calling method^28^, which largely overcame some previous issues such as ‘multimapping’ and ‘exon painting’ in early CAGE datasets used in Faulkner et al. (2009).

We found that the TSSs associated with LTR elements are mainly in the sense strand of LTRs and clustered within narrow regions (**Fig. 2b** for the THE1B subfamily and **Supplementary Fig. 3** for more subfamilies). Since LTR elements contain the promoters for endogenous retroviral elements (ERVs), the sense-biased distributions of TSSs suggest that transcription initiation events in these regions are mainly contributed by the original ERV promoter activities within LTRs. These patterns were not observed in Faulkner et al. (2009), as they only investigated the distributions of TSSs along LTR superfamilies but not the subfamilies. We also found that a large fraction (∼50%) of young TSSs associated with LTRs contain a TATA-box motif starting at 25∼35 bp upstream of the dominant TSSs (**Supplementary Fig. 4**), whereas the ratio drops to ∼30% for the old TSSs associated with LTRs, suggesting a substantial fraction of TATA-box promoters derived from LTRs might have turned into TATA-less promoters during evolution.

**Fig. 3.**
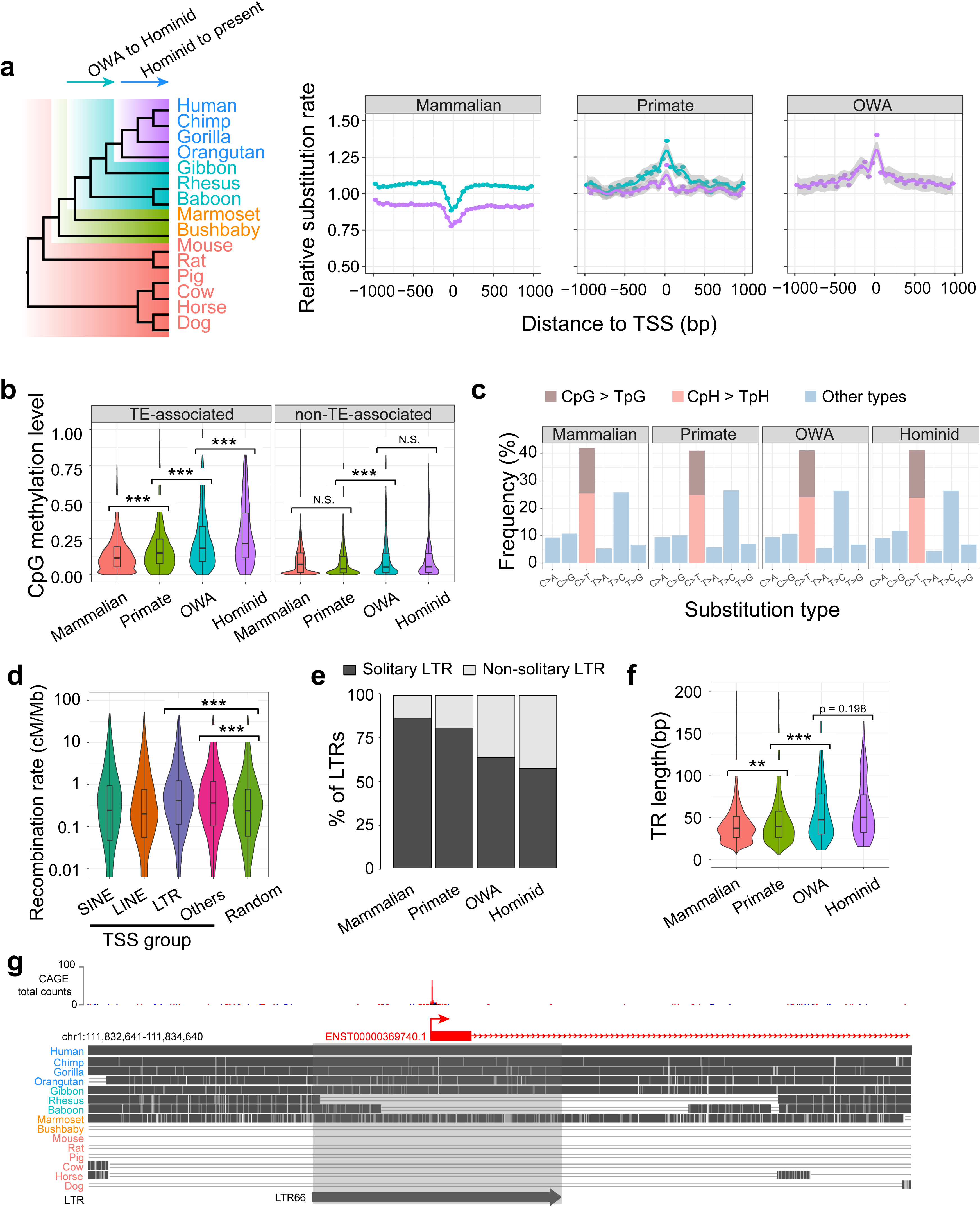
Rapid sequence evolution of young TSSs. (**a**) Left, a phylogeny of genomes used for evolutionary rate analysis, with arrows indicating the two evolutionary periods considered for calculating rates. Right, relative substitution rates (normalized by genomic average) inferred from genomic alignments for three TSS groups, using 40 bins along TSS±1 kb for calculating the average rate in each bin. Best-fit curves were estimated by ‘loess’. (**b**) Violin and box plots for germline DNA methylation levels (a male germline dataset from Guo et al. 2015) for different TSS subgroups defined by the retrotransposon context. For each TSS, the average methylation level of CpGs was calculated for TSS±1 kb. (**c**) Frequencies of nucleotide substitution types in different TSS groups, based on the variants and ancestral alleles from the 1000 genomes project. (**d**) Comparison of recombination rates among TSSs associated with different types of transposable elements and genomic background (‘random’). The recombination rate of each TSS was defined as the average rate for TSS±1 kb. Background recombination rates were generated for randomly selected 2-kb windows in human genome. (**e**) The fraction of solitary LTRs in four TSS groups. (**f**) Distribution of tandem repeat (TR) lengths in four TSS groups. (**g**) An example plot depicting a possible TSS death event around an LTR. Statistical significances in panels **b**, **d** and **f** were calculated by one-tailed Wilcoxon rank sum tests. “*”, p < 0.05; “**”, p < 0.01; “***”, p < 0.001; N.S., not significant.

**Fig. 4.**
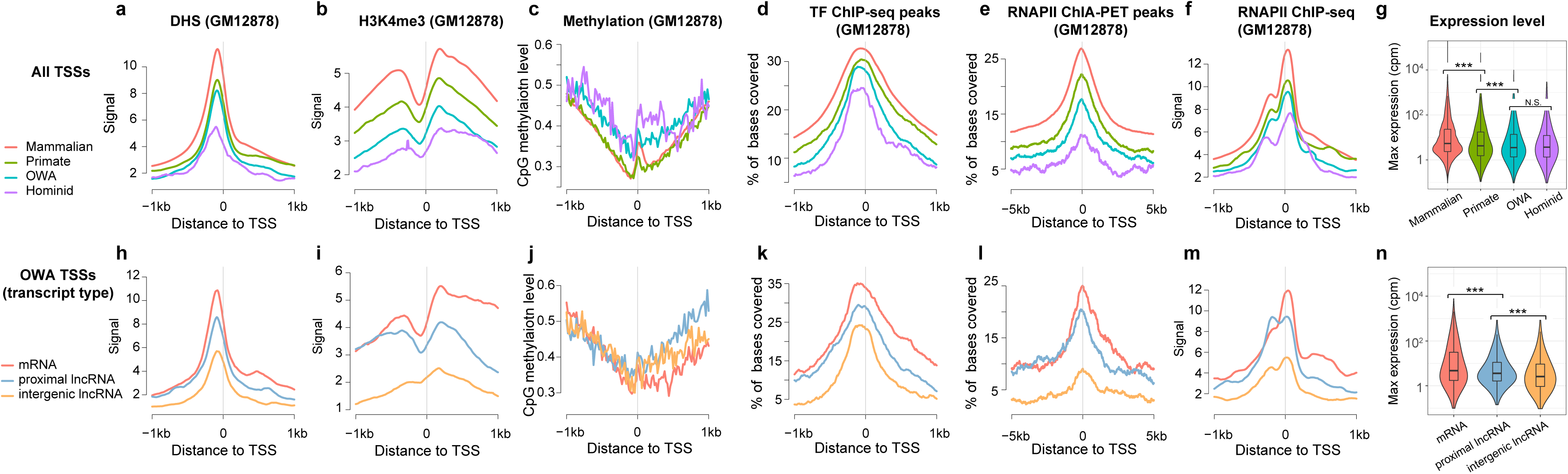
Distinct functional signatures in different TSS groups. (**a**) Meta-profiles of DHS signals for four TSS groups using a 20bp bin size (same bin sizes for other panels). (**b**) Meta-profiles of H3K4me3 signals. (**c**) Meta-profiles of CpG methylation levels. (**d**) Meta-profiles of coverage ratio by TF ChIP-seq peaks. Previously called peaks of 88 TF ChIP-seq datasets from ENCODE were merged together, and for each bin of each TSS locus we calculated how much is covered by merged peaks. (**e**) Meta-profiles of coverage ratio by RNAP II ChIA-PET peaks. (**f**) Meta-profiles of RNAP II ChIP-seq signals. (**g**) Distribution of maximum expression levels of TSSs across primary cell samples, based on the expression data of FANTOM CAT annotation. (**h-n**) Produced using the same methods as for panels **a-g**, but specifically for the OWA TSSs which were divided into subgroups of different transcript types. All functional genomic data except the expression data are for the GM12878 cell line.

LINE-1(L1) is the most abundant LINE family in the human genome (covering ∼20% of human genome). The overall distribution of TSSs along L1 elements (**Fig. 2c**) is similar to that in Faulkner et al. (2009). However, we further observed many differences in the TSS distributions between different L1 subfamilies (**Supplementary Fig. 5**). For some subfamilies, transcription initiation occurs mainly at the region of 5’end antisense promoters (e.g. L1PB1, L1PBa1) which were discussed in Faulkner et al. (2009), whereas for other subfamilies the initiation occurs mainly at the 3’end (e.g. L1MB7) or rather randomly (e.g. L1M4). Although the background distribution of L1 subfamilies in the human genome can explain such difference to some degree, it is apparently not the only reason (**Supplementary Fig. 5)**. This suggests that sequences from different L1 subfamilies have very variable propensity to drive transcription initiation.

**Fig. 5.**
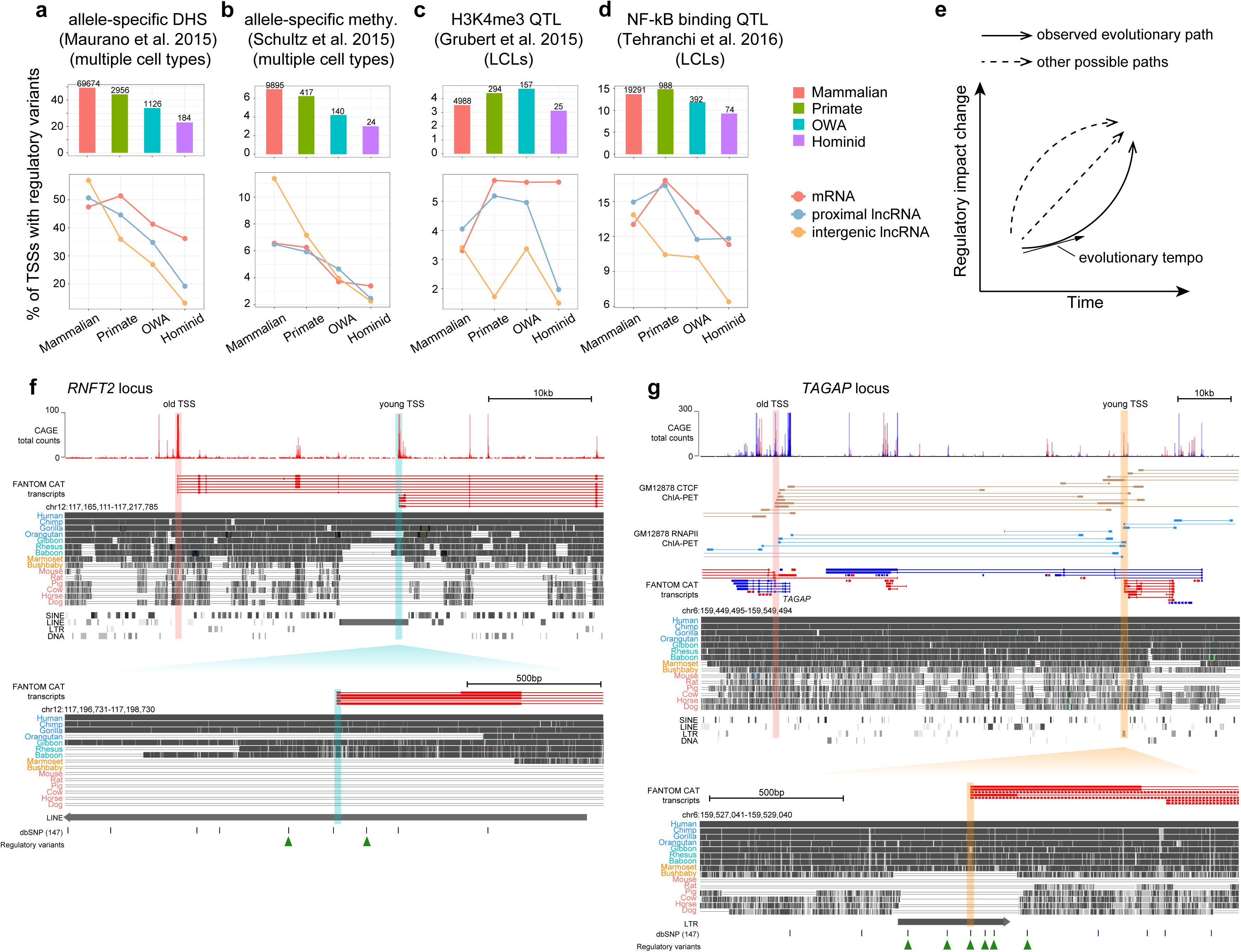
Temporal and spatial constraints on the regulatory evolution of young TSSs. (**a**) Top, proportion of TSSs harboring regulatory variants associated with allele-specific DHS within TSS±1 kb for each TSS group; above the bars are the numbers of TSSs with regulatory variants. Bottom, proportion of TSSs harboring regulatory variants in different TSS subgroups, defined by transcript type. (**b**) Proportion of TSSs harboring variants associated with allele-specific methylation within TSS±1 kb. (**c**) Proportion of TSSs harboring H3K4me3 QTLs within TSS±1 kb. Data generated from lymphoblastoid cell lines (LCLs). (**d**) Proportion of TSSs harboring NF-kb binding QTLs within TSS±1 kb. Data generated from LCLs. (**e**) A schematic illustration depicting different possible evolutionary paths for young TSSs. (**f**) A young TSS *cis*-proximal to old TSSs. Top, FANTOM CAT transcript models (red for forward-strand, blue for reverse-strand); genome alignments and TE annotations obtained from the UCSC genome browser. Bottom, enlarged region of an OWA’ TSS inside a LINE element. Below the alignments are the common SNPs (allele frequency ≥0.01) from the dbSNP database and SNPs associated with regulatory variation within this region. (**g**) A young TSS *trans*-proximal to old TSSs. Top, similar to panel **f** but with additional CTCF and RNAP II ChIA-PET interaction data for GM12878 cell line. Bottom, enlarged region of the young TSS inside a LTR element. Below the alignments are the common SNPs (allele frequency ≥0.01) from dbSNP database and the SNPs associated with regulatory variation within this region.

Alu elements comprise the most abundant SINE family in the human genome (covering ∼10% of human genome). Although Alus are frequently inserted in promoter-proximal and intronic regions, previous research found that they generally lack capacity for driving autonomous transcription^20^. In the FANTOM5 dataset, initially we observed many new TSSs located around the 3’ poly(A) region and the A-rich linker region, but later we found that these TSSs probably resulted from the technical artifacts in the CAGE sequencing in FANTOM5 and thus filtered out the related TSSs (**Supplementary Fig. 6**, see Methods for more details). The remaining Alu-associated TSSs tend to be enriched at the 5’end of Alu in the antisense strand (**Supplementary Fig. 6)**, but how these sequences help drive transcription initiation is unclear.

**Fig. 6.**
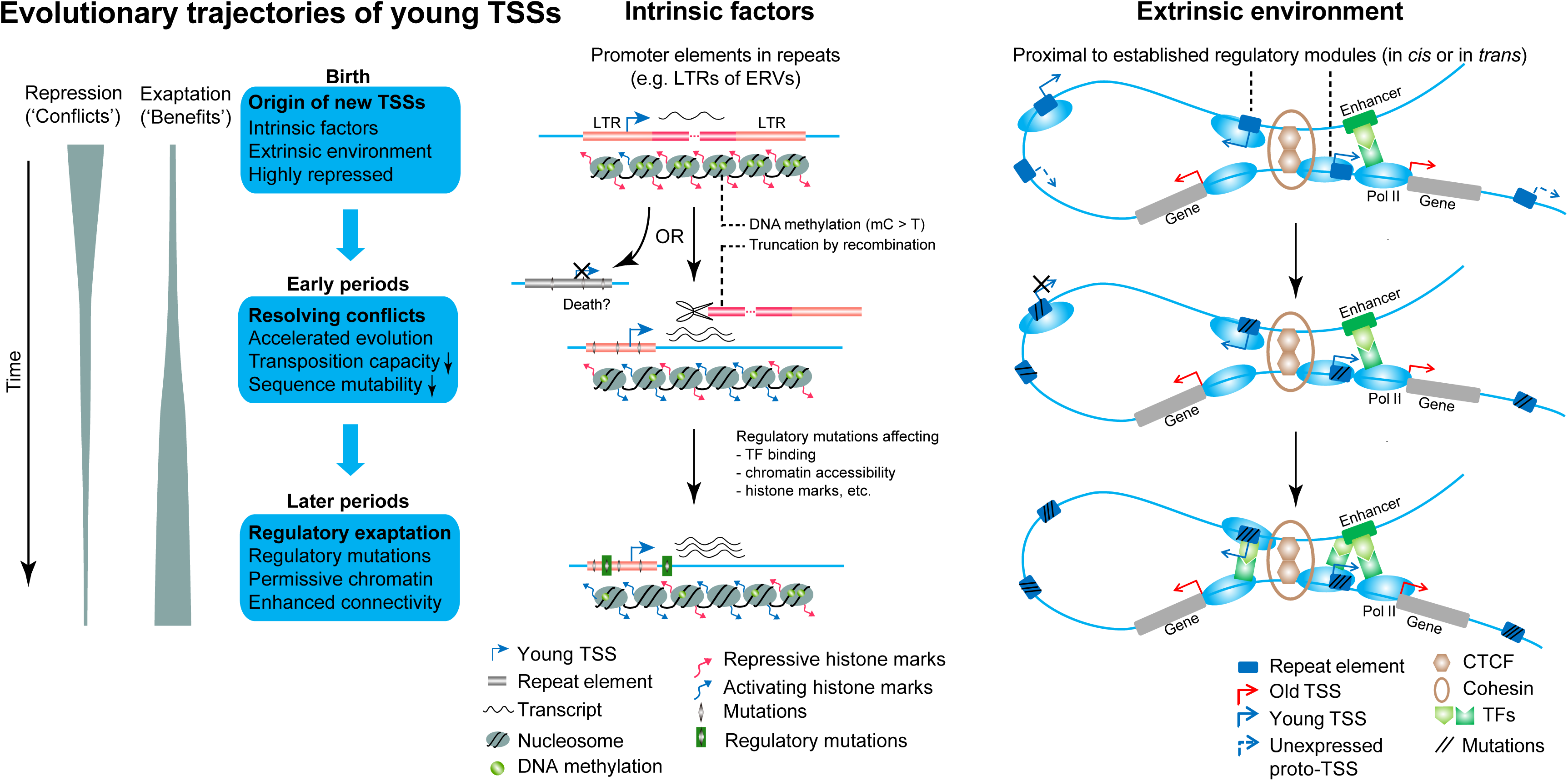
Proposed evolution model for young TSSs. The origin of new TSSs is promoted by sequence-intrinsic and extrinsic factors. A typical intrinsic factor is the promoter element in newly inserted retrotransposons. An important extrinsic factor is the proximity to established regulatory modules as the proximity of a ‘proto-TSS’ to established regulatory elements provides easier access to transcription machinery. Newly emerged TSSs tend to be highly repressed and have limited regulatory capacity. In the early phase, young TSSs undergo rapid sequence evolution allow genomic conflicts associated with repeats to be resolved. Targeted mutational mechanisms enable this rapid evolution, including DNA hypermethylation (methylated C to T mutations), recombination and tandem repeat instability. The accumulated changes around young TSSs can reduce or eliminate the transpositional capacity of associated TEs and stabilize associated tandem repeats. They may also lead to deaths of some young TSSs. In the later phases, surviving TSSs gradually gain mutations in surrounding regions which could increase their regulatory capacity (e.g. TF binding, chromatin accessibility or transcription-associated histone modifications) and are exapted by the host for transcriptional regulation. At the mature phase, TSSs tend to have more permissive chromatin environments, enhanced spatial connectivity and higher expression.

We found that ∼9% of young TSSs contain tandem repeats which are not associated with TEs. Unlike the tandem repeats derived from new TE insertions, the flank regions of these tandem repeats tend to be conserved among mammals and have higher GC content (**Supplementary Fig. 7**), suggesting that some new TSSs in these regions are likely due to autonomous expansions of tandem repeats located in proximal regions of pre-existing promoters (some examples provided in **Supplementary Fig. 7**). This is consistent with previously reported enrichment of tandem repeats in primate promoters^13,22,29,30^.

Taken together, these findings suggest that repetitive sequences significantly contribute to novel TSSs in multiple ways. Among the repetitive sequences, retrotransposons (especially LTRs) are the biggest contributor for generating new TSSs.

#### 2.2.2 Extrinsic factors contribute to novel transcription initiation

Although previous studies and our analyses indicate that some sequence-intrinsic features of repeats can promote transcription initiation, the majority of repeats harboring such proto-TSS sequences do not exhibit initiation signals. For instance, fewer than 1% of LTR elements in the human genome are associated with CAGE-defined TSSs, implying that there are extrinsic factors that could affect the transcription initiation in these regions. One reason for this is that most repeat elements tend to be highly suppressed by the host defense mechanisms, such as DNA methylation and methylation of H3 lysine 9^31^. In addition, we reasoned that proximity of some proto-TSSs to established transcription units might be an extrinsic factor for promoting novel transcription initiation, because such proximity could allow them to access the transcription machinery of other TSSs for initiation. To test this hypothesis, we first examined the *cis*-proximity of the LTR proto-TSSs to old TSSs. Indeed, we found that young TSS-associated LTRs are closer to old TSSs compared to other LTRs that are not associated with TSSs and random genomic intervals (**Fig. 2d**). We further took advantage of published ChIA-PET data which identifies spatially proximal regulatory regions in the genome. We focused on the ChIA-PET data for CTCF and RAD21 (a subunit of cohesin), which are important for chromatin architecture and linking regulatory modules for transcriptional regulation^32^. CTCF binding sites were also found to be highly conserved during evolution^32^. We examined the distances of LTRs to the mammalian-conserved ChIA-PET interaction loci (see Methods) and found that TSS-associated LTRs are closer to CTCF or RAD21 interaction loci compared to non-TSS-associated LTRs (**Fig. 2e**). We suggest that proximity to CTCF/cohesin anchoring loci may enable some proto-TSSs to be spatially proximal to other transcription units and utilize their transcription machinery for initiation.

The spatial proximity of young TSSs to old TSSs may also help to explain the evolution of the number of TSSs per gene. We noticed that the number of TSSs per gene in the human genome approximates to an exponential distribution – the number of genes with a specific number of TSSs decreases exponentially with increase of the number of TSSs per gene (**Fig. 2f**). The exponential relationship appears to be independent of gene lengths, because the it still exists when looking at genes within a specific length range (**Supplementary Fig. 8**). The exponential distribution indicates that most genes have few TSSs, whereas a small fraction of genes have large number of TSSs. A similar relationship is also seen for newly emerged TSSs (**Fig. 2g**), which implies that a small fraction of genes gain many new TSSs during a specific period. We also observed a positive correlation between number of pre-existing TSSs per gene and number of newly gained TSSs per gene (Pearson’s r=0.24, p < 2.2e-16, **Supplementary Fig. 9**) - genes that have more existing TSSs are more likely to gain new TSSs in a later period. Based upon the above observations, we suggest that most of new TSSs derived from repeats arise opportunistically, partly due to their sequence-intrinsic properties and proximity to other transcription units. As time goes by, some newly emerged TSSs could be exapted by proximal genes to form alternative promoters. On the other hand, these observations also suggest that the existing transcriptional landscape to some extent constrains the emergence and evolution of new TSSs.

### 2.3 Rapid sequence evolution of young TSSs

#### 2.3.1 Young TSSs undergo rapid sequence evolution

Next we investigated the subsequent changes of young TSSs after they appear in the genome. One important aspect is the evolutionary rate, which reflects the general trend of sequence evolution. A previous study based on TSSs of early FANTOM projects^14^ showed that evolutionary rates in promoter regions vary between lineages and that the primate lineages appear to have increased rates in promoter regions; however evolutionarily young and old promoters were not separately analyzed. Here we focused on the evolutionary rates for TSS groups of different ages in comparison with the genomic background. To do this, we utilized genomic alignments to infer evolutionary sequence changes around TSS loci for two recent periods (from the last common ancestor of OWAs to the last common ancestor of hominids and from the last common ancestor of hominids to present, as indicated by the phylogeny in **Fig. 3a**), using a maximum likelihood method (see Methods). Based on inferred sequence changes, we calculated the relative rates of substitutions and small insertions/deletions, which were normalized by genomic average. We found that proximal positions of old TSSs have lower substitution rates compared with surrounding regions and genomic average (**Fig. 3a**), suggesting that they were subject to purifying selection in these periods. In contrast, proximal positions of young TSSs exhibit elevated evolutionary rates compared to the surrounding regions as well as genomic average (**Fig. 3a**), suggesting that young TSS loci underwent rapid sequence evolution. Interestingly, for the ‘primate’ TSS group the substitution rates during the early period are higher than in the later period (**Fig. 3a**), suggesting that newly emerged TSSs evolve rapidly at first and then slow down later. Although this pattern is not observed in the insertion/deletion rates (**Supplementary Fig. 10**), it might be due to saturated insertion/deletion mutations and some ancestral insertion/deletion events not being accurately inferred using alignments of extant species. Additionally, by examining the population polymorphism data from the 1000 genomes project, we found that the young TSSs also have elevated variant densities relative to surrounding regions (**Supplementary Fig. 11**), further supporting that young TSSs undergo rapid sequence evolution.

#### 2.3.2 Endogenous mutational processes contribute to rapid evolution of young TSSs

We then asked how the young TSSs evolve rapidly after appearing in the genome. Since many young TSSs are associated with repetitive sequences, we reasoned that some mutational processes associated with repeats could contribute to the rapid evolution.

One contributing factor could be DNA methylation, which is one of main mechanisms for repressing TE activities^31^. We found that the younger TSSs have significantly higher levels of CpG methylation in the germline compared to older TSSs (**Fig. 3b** and **Supplementary Fig. 12)**. In addition, TE-associated TSSs tend to have higher levels of CpG methylation compared to non-TE TSSs within each TSS group (**Fig. 3b**). Because methylated cytosine (mC) can frequently mutate to thymine (T) via deamination, the DNA hypermethylation around young TSSs in the germline represents an important contributor for the elevated evolutionary rates. This is further supported by the substitution patterns in the human population genomic data, in which the C > T is the most common substitution type (∼40% of all substitutions) in all TSS groups and ∼17% of C to T mutations occur in the CpG context (**Fig. 3c**).

Another contributing factor is recombination, which has been found to be associated with mutations and GC-biased gene conversion^33^. We found that LTR-associated TSSs have significantly higher recombination rates relative to genomic average (**Fig. 3d**). Higher recombination rates are also observed in non-TE-associated young TSSs (**Fig. 3d**). Consistently, older LTR-associated TSSs have more solitary LTRs (**Fig. 3e**), which are known to result from allelic or non-allelic homologous recombination^34^. As recombination hotspots evolve rapidly^35^ and ancient recombination events are difficult to detect, it is possible that recombination had also contributed to the rapid evolution of SINE/LINE-associated TSSs.

A third contributing factor is the instability of tandem repeats. Previous research revealed that the mutability of microsatellites (also known as short tandem repeats) increases with their length and long microsatellites tend to be shortened or interrupted by mutations over time^36,37^. Indeed, we found that tandem repeats associated with younger TSSs tend to be shorter than those in older TSSs (**Fig. 3f**), implying that they are more likely to mutate.

#### 2.3.3 Consequences of rapid evolution in young TSSs

A direct consequence of the rapid evolution around young TSSs is that they accumulated many changes, which could reduce or eliminate the transposition capacity of TEs or the mutability of tandem repeats around TSSs, resulting in a more stable genomic environment. Therefore these mutational processes probably help to resolve the genomic conflicts caused by the inherent instability of associated repeats around young TSSs. In addition, we suspect that rapid evolution may lead to deaths of some young TSSs, because some sequence changes could disrupt critical promoter components required for transcription initiation. In the example shown in **Fig. 3g**, a LTR locus with transcription initiation signal in human has been deleted from rhesus and baboon. However, because we lack large-scale CAGE-defined TSSs in other primate species and there could be polymorphisms in TSS loci, we are currently unable to perform detailed analysis regarding the evolutionary deaths of young TSSs.

### 2.4 Functional impact of young TSSs

#### 2.4.1 TSSs of different evolutionary ages exhibit distinct functional signatures

Previous comparison between human and mouse CAGE-defined TSSs revealed that lineage-specific TSSs tend to have tissue-restricted expression profiles, often in samples associated with testis, immunity or brain^13^. Yet how the regulatory functions of these lineage-specific TSSs are gradually established in organisms remain unclear. We sought to investigate the resulting regulatory impact of newly emerged TSSs and how their impact changes over time. We first took advantage of published functional genomic data from ENCODE and other projects to compare related functional signatures between TSS groups, including DNase I hypersensitivity (DHS), histone modifications, DNA methylation, transcription factor (TF) binding and chromatin interactions. Intriguingly, we found that TSSs of different ages exhibit segregating functional signatures (**Fig. 4** for GM12878 cell line) and such patterns are observed in different cell lines (**Supplementary Fig. 13** for K562 and H1-hESC cell lines). Relative to older TSSs, younger TSSs tend to have lower chromatin accessibility (DHS, **Fig. 4a**), lower levels of activating histone modifications (e.g. H3K4me3, H3K27ac, H3K4me1, H3k9ac, **Fig. 4b** and **Supplementary Fig. 14**) and higher CpG methylation (**Fig. 4c**), suggesting younger TSSs are under a more repressed chromatin environment. By examining ChIP-seq data for TFs in ENCODE cell lines, we found that older TSS loci tend to have more binding regions (i.e. more surrounding sequences overlapping ChIP-seq peaks) relative to younger TSSs (**Fig. 4d,** and **Supplementary Fig. 15** for meta-profiles of individual TF ChIP-seq datasets in GM12878). We also observed a similar trend for computationally predicted TFBSs (**Supplementary Fig. 16**). We further analyzed the published ChIA-PET interaction data for RNA polymerase II (RNAP II), which are usually formed within CTCF/cohesin looped structures and considered to reflect promoter-enhancer interactions^38^. We found that younger TSSs have fewer RNAP II chromatin interactions compared with older TSS (**Fig. 4e**), suggesting that younger TSSs tend to lack connections to other regulatory modules. This is consistent with the observations in TF binding (**Fig. 4d**), as TF binding is important for forming promoter-enhancer interactions. As for expression output, younger TSSs tend to display lower expression than older TSSs (**Fig. 4f**), which is consistent with a previous observation that evolutionarily volatile promoters tend to have lower expression levels^13^. Taken together, these observations indicate that the evolution of TSSs leave footprints in the functional signatures of TSSs; namely that younger TSSs tend to have smaller regulatory impact on a genome and that the impact increases with time.

By comparing the TSS subgroups defined by the transcript types, we also observed heterogeneity of functional signatures within TSS groups. Within a similarly-aged group, TSSs associated with mRNAs tend to have higher DHS, more activating histone modifications, more TF binding and more chromatin interactions than other TSSs (**Fig. 4h-m** and **Supplementary Fig. 17**), indicating they are more transcriptionally active. Consistently, mRNA TSSs tend to have higher expression levels than other TSSs within the same group (**Fig. 4n**). Furthermore, TSSs of proximal lncRNAs appear to be more transcriptionally active compared to that of intergenic lncRNAs, likely because they are more proximal to other transcription units. Overall, these findings suggest that locations of young TSSs in gene annotation context could influence the regulatory outcomes.

#### 2.4.2 Evolution of regulatory functions of young TSSs appears to be subject to temporal and spatial constraints

The segregating functional signatures of TSSs of different ages strongly imply that the regulatory outcomes of young TSSs are gradually changed over time. Yet it remains unclear how regulatory changes of young TSSs take place in organisms during evolution, e.g. in what tempo and mode. The regulatory impacts of historical and fixed sequence changes around TSSs are difficult to assess, however, there are many ongoing changes around TSSs within human populations, whose regulatory effects have been widely studied by combining functional and population genomic approaches^39^. Two common strategies are to identify regulatory quantitative trait loci (rQTLs, e.g. TF binding QTLs, histone modification QTLs and DHS QTLs) and variants associated with regulatory allelic specificities (AS, e.g. allele-specific TF binding, allele-specific methylation). Although no QTL or AS study has been specifically performed for human CAGE-defined TSSs, we can apply data from genome-wide rQTL and AS studies of other molecular traits. A previous study^40^ revealed that expression levels of CAGE-defined TSSs are highly correlated with other functional signatures such as TF binding, histone modifications and DHS in surrounding regions, and can be largely predicted by those functional signatures (R^2^ > 0.7). Therefore we reasoned that changes in the regulatory outcomes of TSSs can be approximated by changes in related functional signatures in surrounding regions. By examining rQTLs and AS variants (together called regulatory variants) in TSS loci of different ages, we can gain insights into the tempo and mode of regulatory evolution of TSSs at different life stages.

In our analysis we focused only on the *cis*-regulatory variants around TSS loci, as published *trans*-regulatory variants are rare and of relatively low-quality. Previous expression QTL studies found that the density of *cis*-regulatory variants drops rapidly with increased distances to target TSSs^41^, we restricted our analysis to only regulatory variants within ±1 kb of TSSs. By re-analyzing data from multiple independent studies, including DHS, methylation, histone marks and TF binding, we found that younger TSSs tend to have fewer regulatory variants compared with older TSSs (see **Fig. 5a-d** for four representative datasets and **Supplementary Fig. 18** for more datasets). The trend is especially clear for variants associated with DHS, methylation and TF binding. This is interesting because it suggests that although young TSS loci evolve rapidly, many of the sequence changes appear to have none or limited impact on transcriptional regulation. Since some TSSs are closely spaced, regulatory variants could be counted multiple times in the above analysis (though it may be possible for a variant to affect multiple adjacent TSSs). We still observed similar patterns even after excluding all the TSSs separated by less than 2 kb (**Supplementary Fig. 19**). Moreover, similar trends are observed when only including regulatory variants with high derived allele frequencies (**Supplementary Fig. 20**), changes in which are more likely to be fixed in populations in the future. Overall, these observations imply that regulatory evolution of young TSSs is subject to a temporal constraint - younger TSSs have a slower tempo in regulatory evolution (**Fig. 5e**), which might be due to the strong repression in early periods.

Separating similarly aged TSSs according to transcript type, mRNA and proximal lncRNA TSSs tend to have more regulatory variants compared with intergenic lncRNA TSSs (**Fig. 5a-d**). Since mRNA and proximal lncRNA TSSs also have more ChIA-PET interactions than other TSSs (**Fig. 4l**), we propose that there is a spatial constraint on the regulatory evolution of young TSSs. Generally, younger TSSs have less connectivity to other regulatory modules (i.e. spatially isolated) than older TSSs (**Fig. 4e**), which likely limits their functional impact. In the subsequent evolution, sequence changes in the young TSSs which are proximal to other regulatory modules tend to have more regulatory effects and these TSSs may be incorporated in the existing regulatory network more quickly (i.e. a higher tempo of regulatory evolution in these TSSs). In contrast, relatively isolated TSSs tend to have a slower tempo of regulatory evolution and are more difficult to be co-opted by the host.

Examples of evolving *cis*-proximal and *trans*-proximal young TSSs are shown in **Fig. 5f-g**. In the gene *RNFT2 locus* shown in **Fig. 5f**, an ‘OWA’ TSS, which lies on the antisense strand of a newly inserted L1 element, is *cis*-proximal to an upstream old TSS. In the surrounding regions of the ‘OWA’ TSS, there are multiple polymorphic sites in current populations, two of which are regulatory variants affecting PU.1 binding and H3K4me3 respectively (**Fig. 5f**). In the example shown in **Fig. 5g**, a ‘primate’ TSS within an LTR element is ∼70 kb away from *TAGAP* locus. However, this young TSS is *trans*-proximal to the TSSs of *TAGAP*, as supported by several CTCF and RNAPII ChIA-PET interaction pairs (**Fig. 5g**). This LTR is a solitary LTR and thus lack capacity for retrotransposition. Six regulatory variants are within ±1 kb of the young TSS (**Fig. 5g**). More examples are given in **Supplementary Fig. 21**.

## 3 Discussion

Given the large number of identified TSSs in the mammalian genomes and the high TSS turnover rate, it is important to understand where the new TSSs come from, how they evolve over time, and their functional impact on transcripts. By performing evolutionary and functional analyses, we gain several important insights into the evolution of newly emerged TSSs. We summarize our main findings in an integrative model as shown in **Fig. 6**.

First, our analyses revealed several sequence-intrinsic and extrinsic factors that promote the emergence of new TSSs (**Fig. 6**). Intrinsic factors are mainly associated with the expansion of repetitive sequences, among which retrotransposons represent a major source of new TSSs. In addition to sequence-intrinsic properties, chromatin organization and spatial chromosomal interactions are likely important extrinsic factors. New TSSs are usually proximal in *cis* or *trans* to other established transcriptional units providing easier access to the transcriptional machinery, whereas unexpressed proto-TSSs are more isolated. This dependence on extrinsic chromatin environment partly explains why only a small fraction of proto-TSSs have detectable initiation signals.

Secondly, resolving genomic conflicts is likely the main theme in the early period of young TSSs (**Fig. 6**). Our evolutionary rate analysis revealed that young TSSs experienced rapid sequence evolution in early periods, which appear to be associated with several endogenous mutational processes, including DNA methylation, recombination and tandem repeat mutagenesis. We suggest that such rapid evolution can reduce the genomic conflicts caused by the instability of repetitive sequences associated with young TSSs, as the TSS loci became more stable after they mutated. We suspect that a considerable fraction of new TSSs may die during the rapid evolution in early periods, as sequence changes could disrupt critical promoter components required for transcription initiation.

Thirdly, by analyzing functional genomic data, we found that in early periods young TSSs tend to have limited transcriptional competency, likely due to the highly repressive environment and lack of connectivity to other functional modules. However, their regulatory potential appear to be gradually enhanced over time (**Fig. 6**). Interestingly, by examining regulatory variants around TSS loci, we revealed that the evolution of regulatory functions of young TSSs appears to be subject to temporal and spatial constraints. The temporal constraint - that younger TSSs have fewer regulatory variants within a period (slower tempo) despite faster sequence evolution - is probably due to the genomic conflicts caused by the novel transcription and associated unstable repetitive sequences. Young TSSs tend to be strongly repressed at first and require time to resolve the genomic conflicts caused by associated repeats. The spatial constraint – that TSSs with fewer chromosomal contact display a slower tempo of regulatory evolution - likely limits the regulatory impact of young TSSs in early stages and affects the evolutionary trajectories of young TSSs depending on their genomic context. Based upon these observations and proposed constraints, we speculate that younger and (or) more isolated TSSs are more likely to die out during evolution.

Many studies have reported the contribution of repetitive sequences to regulatory innovation^34^. Our detailed analysis on evolutionary trajectories of young human TSSs provide new strong evidence. We have shown that the repeat-derived TSSs are tightly constrained in the beginning and have limited functional impact, but after resolving genomic conflicts some are successfully incorporated into the existing regulatory network, turning “conflicts” into “benefits”^34^. In the long run, the repeat-derived TSSs contribute significantly to regulatory innovation. Interestingly, a similar evolutionary pattern was also observed in Alu exonization in primate genomes^42^, implying a commonly used strategy in genome evolution. Given the pervasiveness of repetitive sequences and the similarity of chromatin structures in eukaryotic genomes, the observed evolutionary processes involved in newly emerged TSSs in primate genomes could also exist in other eukaryotic groups. These evolutionary patterns also suggest the importance of balancing evolvability and robustness in genome evolution^43^.

## Methods

### Human TSS annotation dataset

We used the FANTOM 5 TSS dataset because it is the most comprehensive TSS annotation to date, cataloguing/encompassing the genome-wide TSS profiling of most major primary cell types and tissues in human. The high-confidence, “robust” TSSs from the latest FANTOM CAT annotation (http://fantom.gsc.riken.jp/cat/, part of FANTOM 5)^28^ were used for our analyses, particularly as each TSS has been assigned a RNA-seq-defined transcript. Coding status and transcript classification of transcripts were defined as in the FANTOM CAT. To facilitate analysis and interpretation, we merged three lncRNA classes (“lncRNA_antisense”, “lncRNA_divergent” and “lncRNA_sense_intronic”) in the FANTOM CAT annotation into a class called “proximal lncRNA”, because these lncRNAs are proximal to other transcript units. We also merged several minor classes (“sense_overlap_RNA”, “short_ncRNA”, “small_RNA”, “structural_RNA” and “uncertain_coding”) into a class called “other RNA”. For TSSs which are associated with multiple types of transcripts, we assigned them hierarchically to the five categories: mRNA > proximal_lncRNA > intergenic_lncRNA > pseudogene > other_RNA. As CAGE TSS peaks (i.e. tag clusters) usually span more than 1 bp, unless specified otherwise, we used the dominant TSS position (i.e. the most frequently used initiation site) of each TSS peak provided in the FANTOM annotation for most analyses.

### Categorization of human TSSs by sequence age

To categorize human TSSs by the evolutionary age of the sequence, we made use of whole genome alignments between human (hg19) and 16 other mammalian genomes (**Supplementary Table 1**) from UCSC genome browser^44^. To estimate the sequence ages of human TSS loci, the UCSC liftOver tool was used to determine presence or absence of each human TSS sequence in other non-human genomes based on available pairwise chain alignment files from UCSC. We required a minimum mapping ratio of 90% for CAGE TSS peaks (∼23bp in length on average), which usually covers Initiator (Inr) elements of promoters. The sequence proximal to Inr element has previously been found to be conserved in mammalian promoters^14^. In addition, we required a minimum mapping ratio of 50% for TSS peaks±100 bp, which we considered as “core promoter” regions in our study and are usually under high selective constraint^14^, although there is no standard definition for “core promoter” currently. To reduce potential false positives resulting from alignments of paralogous loci in two genomes, we further required a minimum alignment chain size of 10 kb for both target and query genomes. A human TSS locus satisfying the above criteria for the pairwise alignment was considered as having the orthologous sequence in the surveyed genome, and its sequence age should be equal to or larger than the age of last common ancestor of two species. The presence/absence patterns of TSSs were then used for defining the four TSS groups as described in the main text. We also tried multiple sets of thresholds for liftOver which did not result in notable variation in the grouping results (**Supplementary Table 2**), mainly because many newly emerged TSS loci were associated with TE insertions, which usually span more than 200 bp.

As some genomic regions are highly repetitive and could lead to poor assemblies and erroneous alignments, we filtered out any TSS whose ±1 kb regions overlapping the blacklisted genomic regions (see **Supplementary Table 3**) defined in the ENCODE project and two other studies^45,46^. Because CAGE reads are usually short (20∼70bp)^27^ and can be mapped to the genome multiple times, we made use of the Duke 20-bp uniqueness track from UCSC browser to filter out the TSS peaks that have an average uniqueness score of <0.5 (a 20-bp uniqueness score of <0.5 means that a 20-mer can be mapped to the human genome more than twice). After excluding these blacklist regions, we still observed that some TSS loci, which are usually associated with low-complexity tandem repeats, exhibited suspiciously high read depths in some functional datasets, suggesting they might be artifacts due to poor mappability for short reads in those regions. Therefore we further filtered out any TSS harboring more than 10% (200 bp) of tandem repeats in the 2 kb region centered on the TSS. In addition, TSSs of chrM and chrY were excluded from all analyses because some genome assemblies or functional datasets lack data for these genomic sequences.

When analyzing the remaining TSSs, we further found two significant sources of putative false positives. One is the pseudogene-associated TSSs. Pseudogenes (especially processed pseudogenes) were reported as a notable source of false positives for CAGE-defined TSSs because of their high sequence similarity to original gene loci and the short lengths of CAGE reads^47^. For the GM12878 cell line, only 3.7% of the pseudogene TSSs in primate lineages from FANTOM 5 can be found in the previously published GRO-cap-defined TSSs (**Supplementary Fig. 6**)^5^.

Therefore we excluded all pseudogene TSSs from downstream analyses. Another source of false positives is the TSSs associated with poly(A) or poly(T) tracts. We initially found many young TSSs in FANTOM 5 located around the 3’ poly(A) region and the A-rich linker region of Alu elements. However, in the GM12878 cell line, only 5.2% of the poly(dA:dT)-associated TSSs in primate lineages from FANTOM 5 can be found in the GRO-cap-defined TSSs (**Supplementary Fig. 6**). On the other hand, a much larger fraction (43%) of the TSSs that are not associated with pseudogenes and poly(dA:dT) tracts can be found in the GRO-cap-defined TSS dataset. Such a large difference in the overlapping ratio suggests that the TSSs associated with poly(dA:dT) tracts have a high fraction of false positives. A recent study also suggested that Alu sequences generally lack the capacity to drive autonomous transcription^20^. Therefore we filtered out the TSSs flanked by a tandem repeat with A content of >50 % or T content of >50 % within ±100 bp.

### Analysis of TATA-box and CpG islands (CGI)

The data of CGI annotation in the human genome was from Cohen et al. (2011)^48^. A TSS was considered as CGI-associated if its core promoter region (TSS±100 bp) overlaps a CGI. TATA-box hits were predicted by R package “seqPattern” using the TBP position-weighted matrix with a minimum score of 80%. A TSS was considered as TATA-box-associated if the start of a TATA-box motif is located at 25∼35 bp upstream of the TSS.

### Analysis of repeats associated with TSSs

The annotation of transposable elements in our analysis was based on RepeatMasker annotation of the hg19 assembly, downloaded from http://www.repeatmasker.org (Repeat Library 20140131)^24^. In addition, as young TSS loci are frequently associated with tandem repeats, tandem repeats annotated by TRF (downloaded from UCSC) and STRcat^26^ were also used. The “Simple repeat”, “Low complexity” and “Satellite” families in RepeatMasker were considered as tandem repeats in our analysis. The tandem repeats from RepeatMasker, TRF and STRcat were merged into a union dataset. For overlapping tandem repeats in these three datasets, the priority order for being included in the union dataset was STRcat > TRF > RepeatMasker.

To investigate the repeat content around TSS loci, we first identified the nearest repeat element to each TSS and counted how many TSSs harbored repeat elements within TSS±100 bp regions (i.e. core promoter regions in this study). Since retrotransposons and tandem repeats were the main types of TSS-associated repeats and many tandem repeats were derived from retrotransposons, for each TSS group defined by sequence age, we further defined four TSS subgroups (‘SINE-associated’, ‘LINE-associated’, ‘LTR-associated’ and ‘Others’) based on the nearest retrotransposon within 100 bp of the TSS. The statistics of subgroups defined by transcript types and associated retrotransposons are given in **Supplementary Table 4.**

To analyze the distributions of TSSs along repeat elements, we calculated the relative distances of TSSs to the 5’ (corresponding to 0% of the full-length) of corresponding repeat subfamily consensus sequences based on the alignment information provided in RepeatMasker annotation. When investigating distances of young TSSs to ChIA-PET interaction loci of CTCF or RAD21, we only considered the interaction pairs whose sequences could be found in at least one of the six non-primate mammalian genomes listed in **Supplementary Table 1**, based on the liftOver mapping with parameters “-minMatch=0.5 -minChainT=10000 -minChainQ=10000”. The chromatin interactions in these mammalian-conserved loci are likely established before emergence of primates and conserved among mammals.

### Evolutionary rate analysis

To investigate the evolutionary rates around TSS loci, we extracted alignments of human and 14 other mammalian genomes for all TSS and their surrounding 2 kb regions from the 100-way MULTIZ genome alignments from UCSC (all species used for analysis are listed in **Supplementary Table 1**; tarSyr1 and micMur1 were not in the 100-way alignments and thus not included in this analysis). To improve the alignment quality, the extracted MULTIZ alignments were re-aligned using PRANK with parameter “+F”, which was found to generate more accurate gapped alignments for evolutionary analysis ^49^. The re-alignment results were then used to infer ancestral sequences for each TSS locus using FASTML^50^ with parameters “--SubMatrix HKY -jointReconstruction no --indelReconstruction ML”. FASTML produced posterior probabilities for each position of inferred ancestral sequences. Positions with low-confidence inferred sequences (maximum marginal probability of <0.8) were excluded for subsequent analyses. Evolutionary sequence changes (substitutions, insertions and deletions) in TSS loci in different periods were identified by comparing inferred ancestral sequences and derived sequences, and these changes were used to calculate substitution, insertion, and deletion rates for each period respectively. To estimate the genomic average evolutionary rates, we generated 10,000 random 2-kb intervals from the human genome, and ran the same analysis pipeline as described above for the TSS loci. The relative rates of substitutions, insertions and deletions in TSS loci were then obtained by dividing the original rates by genomic average rates estimated from random intervals.

### Analysis of mutational mechanisms

Because spontaneous deamination of methylated cytosines (causing cytosine to thymine substitutions) was found to be a major source of mutations during evolution, we analyzed the germline DNA methylation levels to investigate the impact of methylation on the evolutionary rates of different TSS groups. We used the published germline DNA methylation data from Guo et al. (2015)^51^ and focused on the CpG methylation events. The methylome of male primordial germ cells of 7-weeks old embryos was used in our analysis, because this sample exhibited a high degree of methylation across the genome, as shown in that study. Data of recombination rates in human populations was from the HapMap project^52^. The completeness status (solitary or non-solitary) of LTRs was predicted by REannotate^53^ with parameters “-n -c”, using the RepeatMasker annotation as input.

### Analysis of functional signatures of TSSs

Processed data (files of normalized signals and called peaks) of Dnase I-seq, ChIP-seq and DNA methylation (WGBS) of ENCODE cell lines (GM12878, K562 and H1-hESC) were downloaded from ENCODE website and ENSEMBL database. Analysis and visualization of functional genomics data on the TSS groups and subgroups were performed with BEDtools, R, seqplots^54^ and deeptools^55^.

ChIA-PET data for CTCF and RNAPII in GM12878 were from Tang et al. (2015)^38^. ChIA-PET data for RAD21 in GM12878 were from Grubert et al. (2015)^56^. ChIA-PET data for RNAPII in K562 cell line were downloaded from the ENCODE website.

### Regulatory variant analysis

The ongoing genomic changes (polymorphic sites) affecting the regulatory outcomes of TSSs in human populations can be considered as a snapshot of regulatory evolution of TSSs. Investigation of these regulatory variants would help to understand how the regulatory impact of TSSs changes over time. Therefore, we analyzed published regulatory variants which affect transcription-related molecular traits, such as TF binding, histone marks, DNA methylation and DNase I hypersensitivity (DHS) from several genome-wide studies.

Regulatory variants for allele-specific DHS in multiple cell types were from Maurano et al. (2015)^57^. Regulatory variants for allele-specific CpG methylation in multiple cell types were from Schultz et al. (2015)^58^. Three types of histone mark QTLs (H3K4me3, H3K4me1 and H3K27ac) of lymphoblastoid cell lines (LCLs) were from Grubert et al. (2015)^56^. For the data from Grubert et al. (2015), we only used the regulatory variants that are located within the corresponding regulated histone peak regions for analysis. Binding QTLs of 5 TFs (JunD, NF-kb, Pou2f1, PU.1 and Stat1) and H3K4me3 QTLs in LCLs were from Tehranchi et al. (2016)^59^. The derived allele frequencies (DAFs) of variants were based on the data of 1000 genomes project phase 3 release and only variants with known ancestral alleles were used for analysis. For each type of regulatory variant, we calculated the proportion of TSSs harboring at least one regulatory variant within 1 kb of the TSS. To account for the issue of possible duplicated counts of adjacent TSS loci, we repeated the analysis after excluding all the TSSs separated by less than 2 kb for to prevent duplicated counts. We also repeated the analysis for datasets under three different minimum DAFs (0.01, 0.1 and 0.5).

## Data availability

All the analyses in this study were based on published datasets. A table of data source links is given in **Supplementary Table 5**. A table containing the defined TSS groups/subgroups in this study is provided in **Supplementary Table 6.** All other data are available from the authors upon reasonable request.

## Acknowledgments

We are most grateful to Jernej Ule, Anna Poetsch, Jan Attig and Anob Chakrabarti for insightful comments on earlier versions of the manuscript. We also thank all the members of Luscombe lab for helpful advice and discussions throughout the project. This work is supported by the Francis Crick Institute which receives its core funding from Cancer Research UK (FC001110), the UK Medical Research Council (FC001110), and the Wellcome Trust (FC001110) (N.M.L.). N.M.L. is also supported by a Wellcome Trust Investigator Award and core funding from the Okinawa Institute of Science & Technology. C.L. is funded by a EMBO long-term postdoctoral fellowship (ALTF 1499-2016).

## Author Contributions

C.L. conceived the project, with considerable discussion with N.M.L. C.L. performed the analyses and drafted the manuscript; N.M.L. supervised the project and contributed extensively to the writing and revising of the manuscript. B. L. provided important advice and contributed to the writing and revising of the manuscript.

## Competing financial interests

The authors declare no competing financial interests.

## Supplementary Information

**Table S1.**
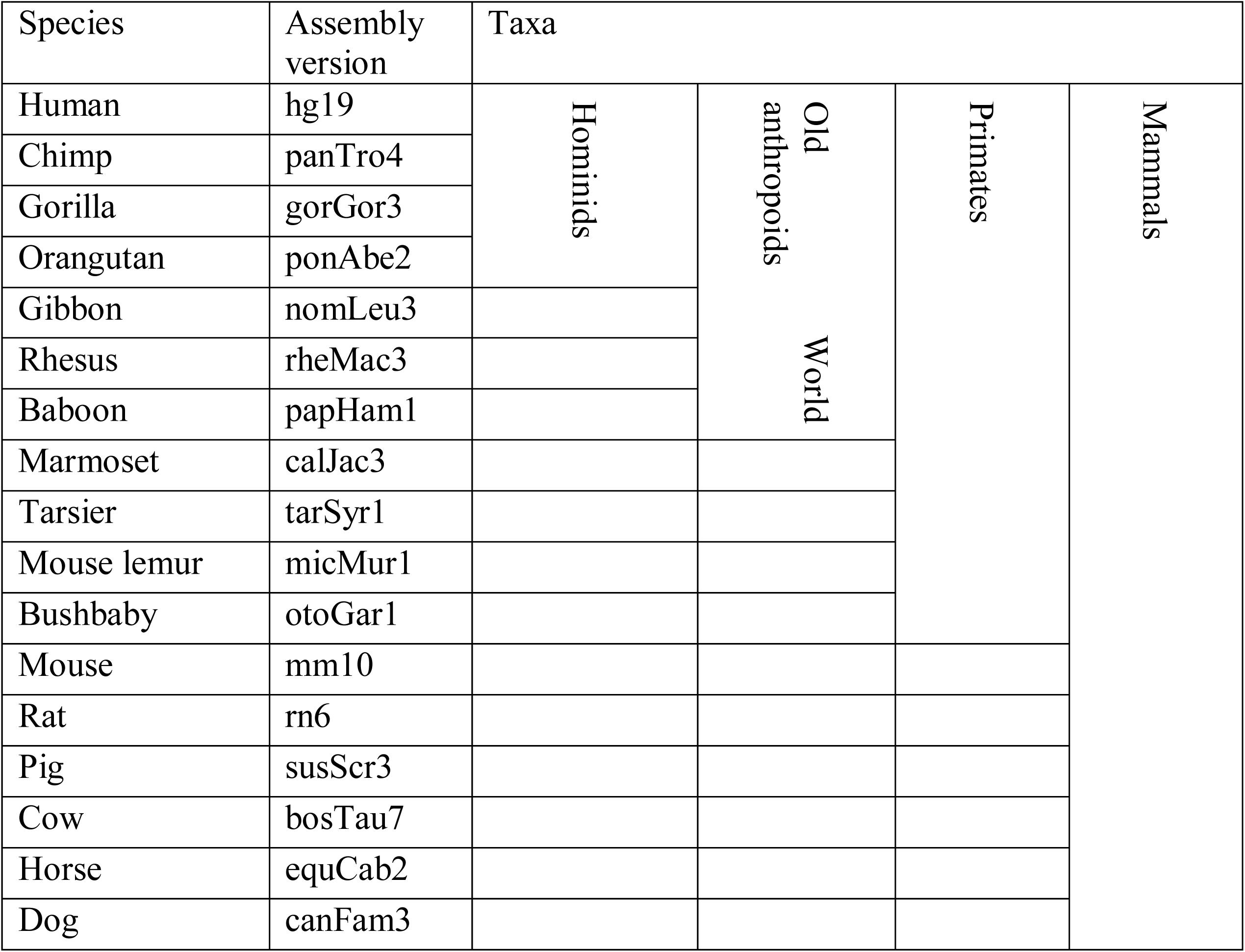
Species and genome assemblies used for estimating sequence ages of TSSs.

**Table S2.**
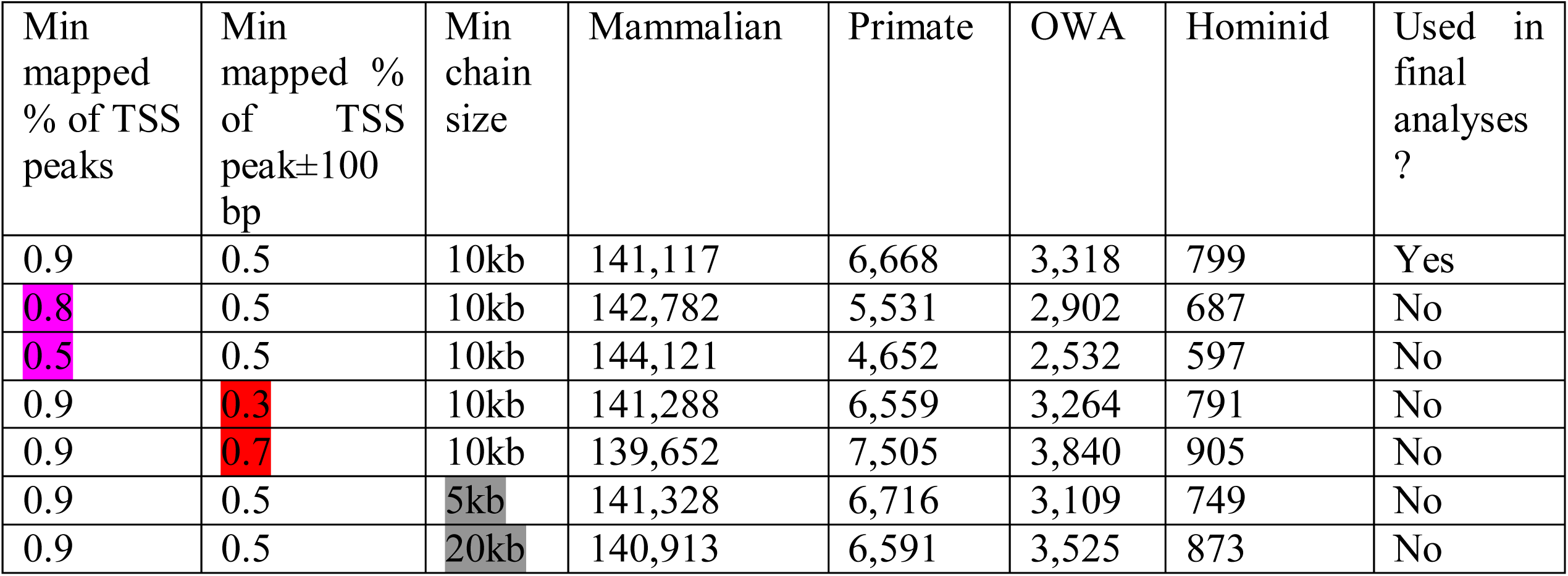
Statistics of grouping results with different sets of cutoffs for liftOver, after filtering the TSSs overlapping blacklist regions.

| Min mapped % of TSS peaks | Min mapped % of TSS peak±100 bp | Min chain size | Mammalian | Primate | OWA | Hominid | Used in final analyses ? |
| --- | --- | --- | --- | --- | --- | --- | --- |
| 0.9 | 0.5 | 10kb | 141,117 | 6,668 | 3,318 | 799 | Yes |
| 0.8 | 0.5 | 10kb | 142,782 | 5,531 | 2,902 | 687 | No |
| 0.5 | 0.5 | 10kb | 144,121 | 4,652 | 2,532 | 597 | No |
| 0.9 | 0.3 | 10kb | 141,288 | 6,559 | 3,264 | 791 | No |
| 0.9 | 0.7 | 10kb | 139,652 | 7,505 | 3,840 | 905 | No |
| 0.9 | 0.5 | 5kb | 141,328 | 6,716 | 3,109 | 749 | No |
| 0.9 | 0.5 | 20kb | 140,913 | 6,591 | 3,525 | 873 | No |

**Table S3.** Lists of blacklist genomic regions used for filtering TSSs.

| File | Source |
| --- | --- |
| wgEncodeDukeMapabilityRegionsExcludable.bed | ENCODE |
| wgEncodeDacMapabilityConsensusExcludable.bed | ENCODE |
| seq.cov1.ONHG19.bed | Pickrell et al. 2011 |
| UM1K0M50BP.bed | Li and Freudenberg 2014 |

**Table S4.** Statistics of TSS subgroups defined by transcript types and the nearest retrotransposon elements.

| Mammalian |  |  | Primate |  |  | Old World Anthropoid |  |  | Hominid |  |  |
| --- | --- | --- | --- | --- | --- | --- | --- | --- | --- | --- | --- |
| mRNA | SINE | 3427 | mRNA | SINE | 271 | mRNA | SINE | 100 | mRNA | SINE | 16 |
|  | LINE | 3470 |  | LINE | 299 |  | LINE | 118 |  | LINE | 43 |
|  | LTR | 830 |  | LTR | 433 |  | LTR | 306 |  | LTR | 61 |
|  | Others | 75301 |  | Others | 1569 |  | Others | 555 |  | Others | 145 |
| proximal lncRNA | SINE | 2019 | proximal lncRNA | SINE | 204 | proximal lncRNA | SINE | 89 | proximal lncRNA | SINE | 15 |
|  | LINE | 2311 |  | LINE | 266 |  | LINE | 164 |  | LINE | 34 |
|  | LTR | 827 |  | LTR | 465 |  | LTR | 309 |  | LTR | 67 |
|  | Others | 35202 |  | Others | 1071 |  | Others | 426 |  | Others | 87 |
| intergenic lncRNA | SINE | 966 | intergenic lncRNA | SINE | 84 | intergenic lncRNA | SINE | 43 | intergenic lncRNA | SINE | 8 |
|  | LINE | 1232 |  | LINE | 219 |  | LINE | 173 |  | LINE | 53 |
|  | LTR | 1192 |  | LTR | 799 |  | LTR | 524 |  | LTR | 146 |
|  | Others | 9106 |  | Others | 516 |  | Others | 269 |  | Others | 58 |
| other RNA | SINE | 324 | other RNA | SINE | 32 | other RNA | SINE | 11 | other RNA | SINE | 0 |
|  | LINE | 368 |  | LINE | 78 |  | LINE | 39 |  | LINE | 17 |
|  | LTR | 147 |  | LTR | 99 |  | LTR | 67 |  | LTR | 17 |
|  | Others | 4395 |  | Others | 263 |  | Others | 125 |  | Others | 32 |

**Table S5.**
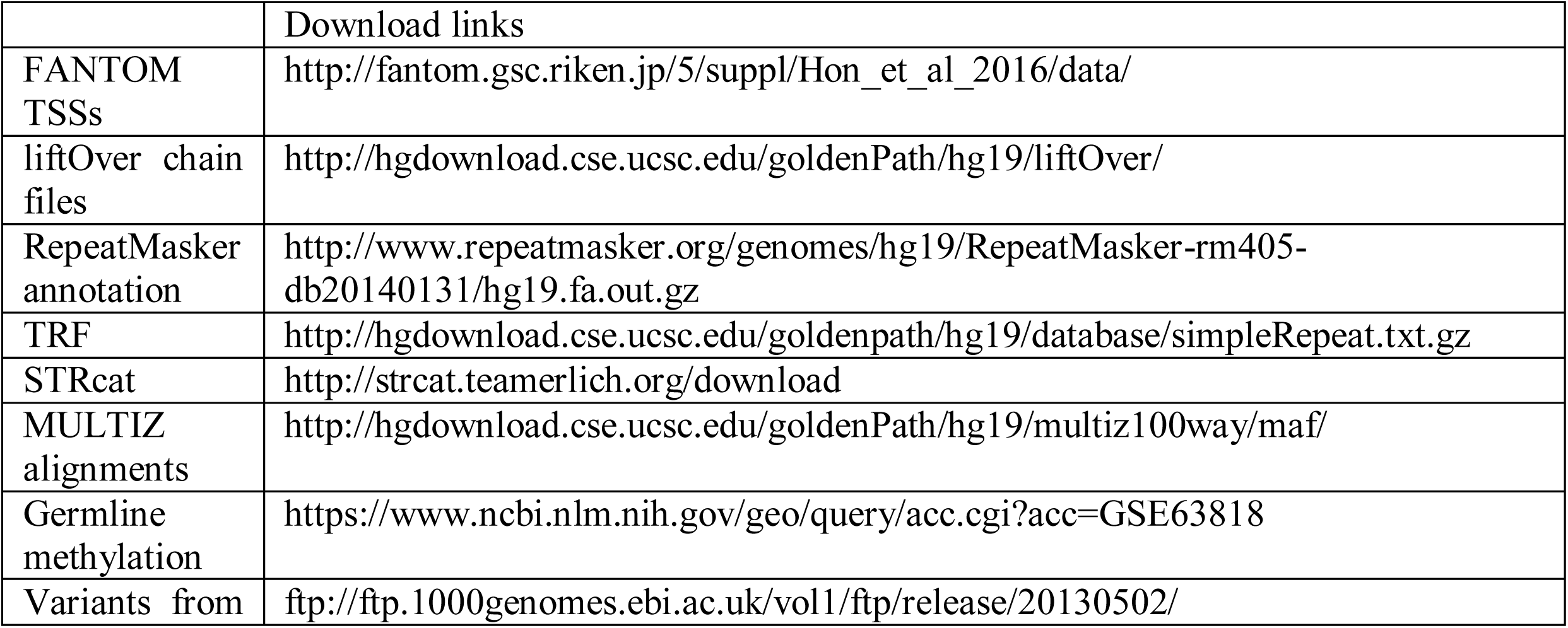
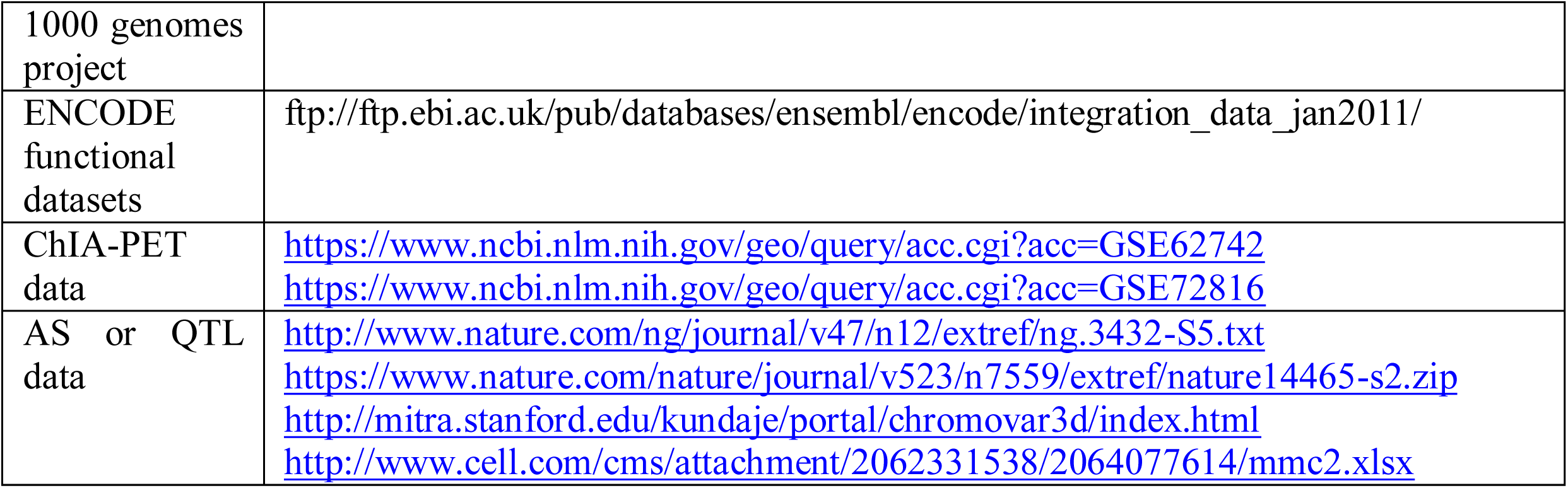
URL links of main published datasets used in this study.

**Supplementary Table 6 A table containing the defined TSS groups/subgroups used in analyses (in a separate file).**

**Figure S1.**
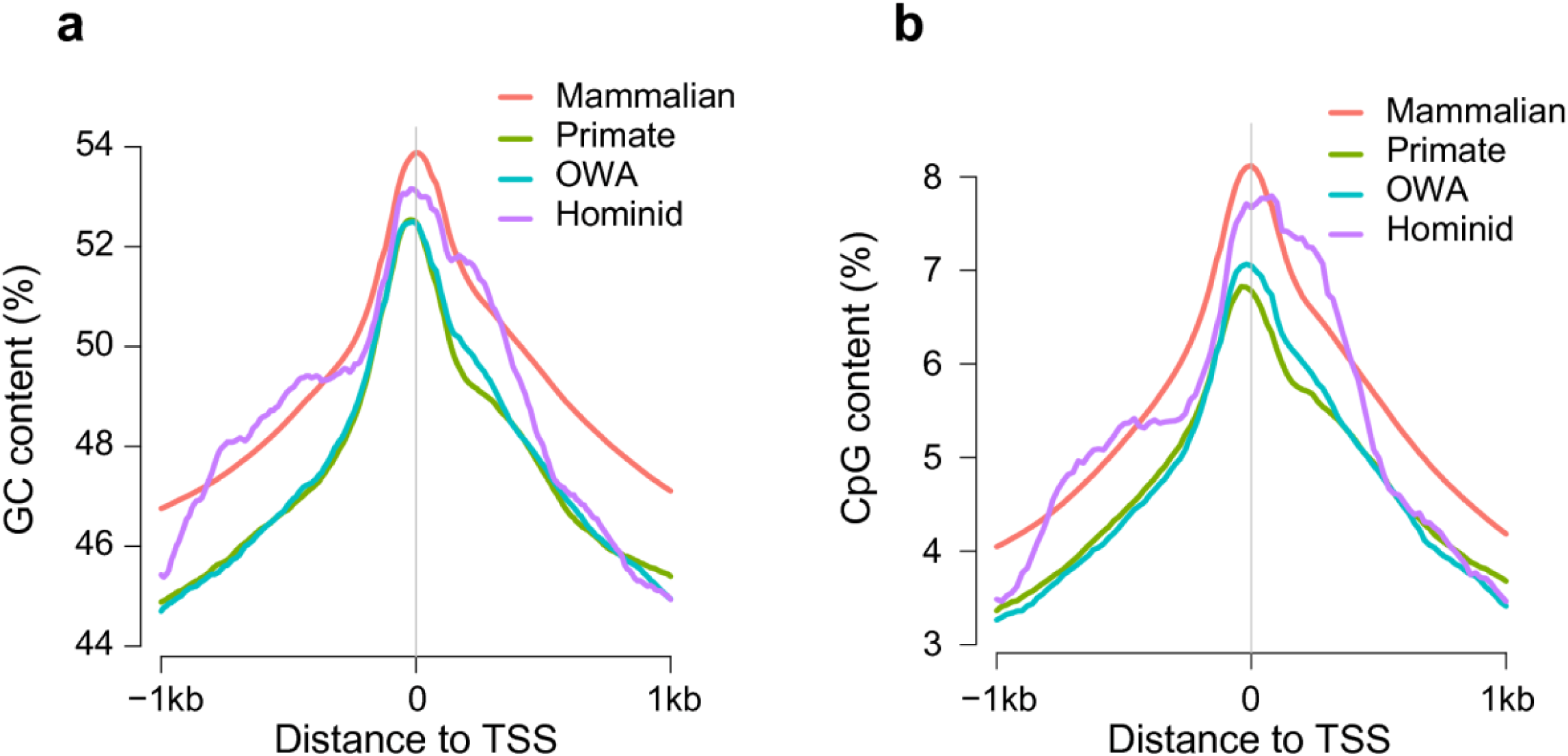
Comparison of GC content and CpG content between four groups.

**Figure S2.**
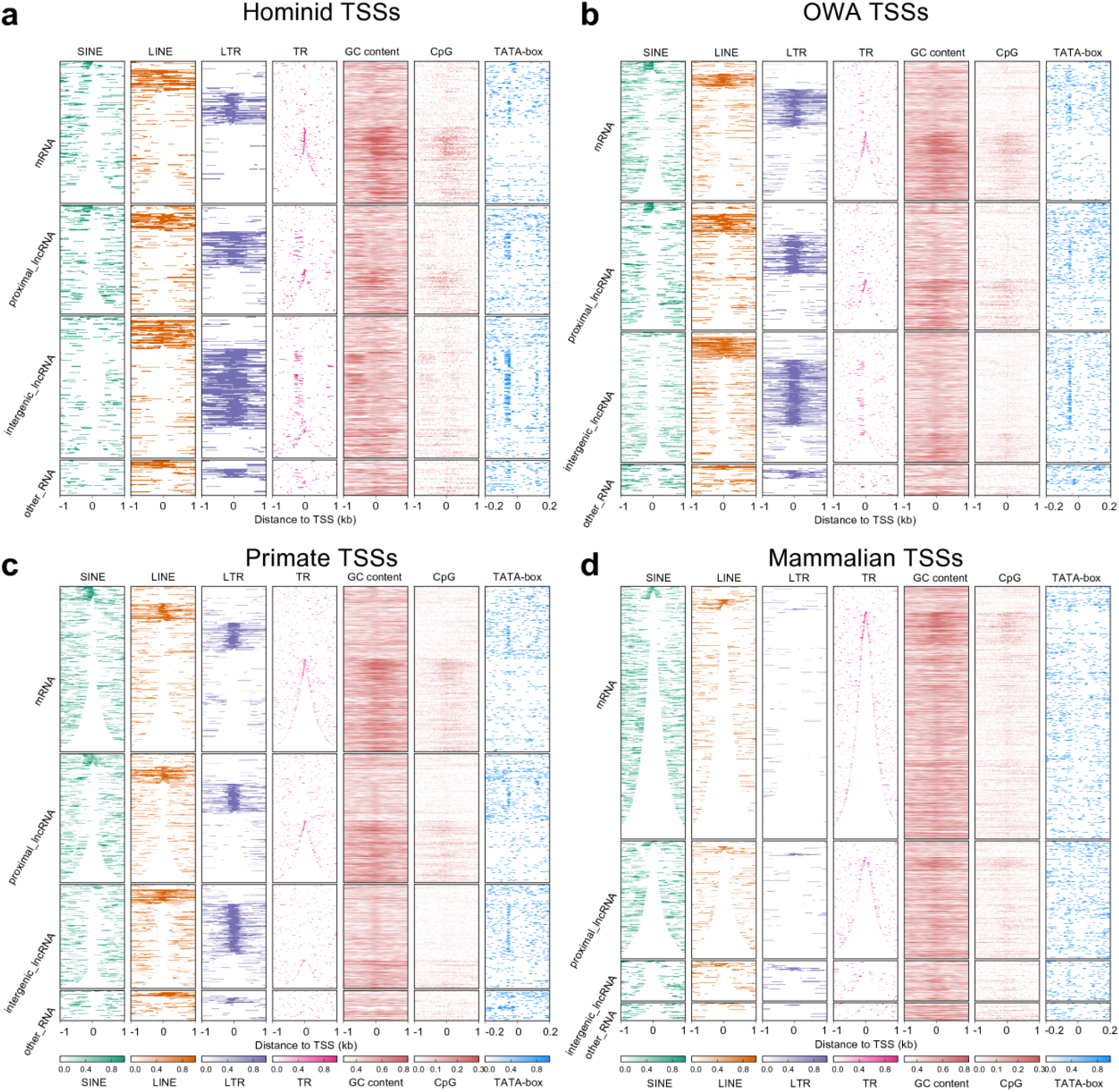
Heatmap for repeat content, GC content, CpG content and TATA-box in four TSS groups. Each TSS group is subdivided into subgroups based on transcript type. Within each subgroup, rows are sorted by the distance from the TSS to the nearest TE element (priority order: SINE > LINE > LTR > Others). For the TSSs in the ‘Others’ category, rows are sorted based on their distances to the nearest tandem repeat elements. The color gradients for repeat elements (SINE, LINE, LTR, tandem repeat (TR)) represent the repeat coverage in 10 bp bins.Regions shown in the TATA-box columns are TSS±200 bp. Because of the large number of TSSs in the ‘mammalian’ group (panel **d**), only the data of 5000 randomly selected TSSs are shown.

**Figure S3.**
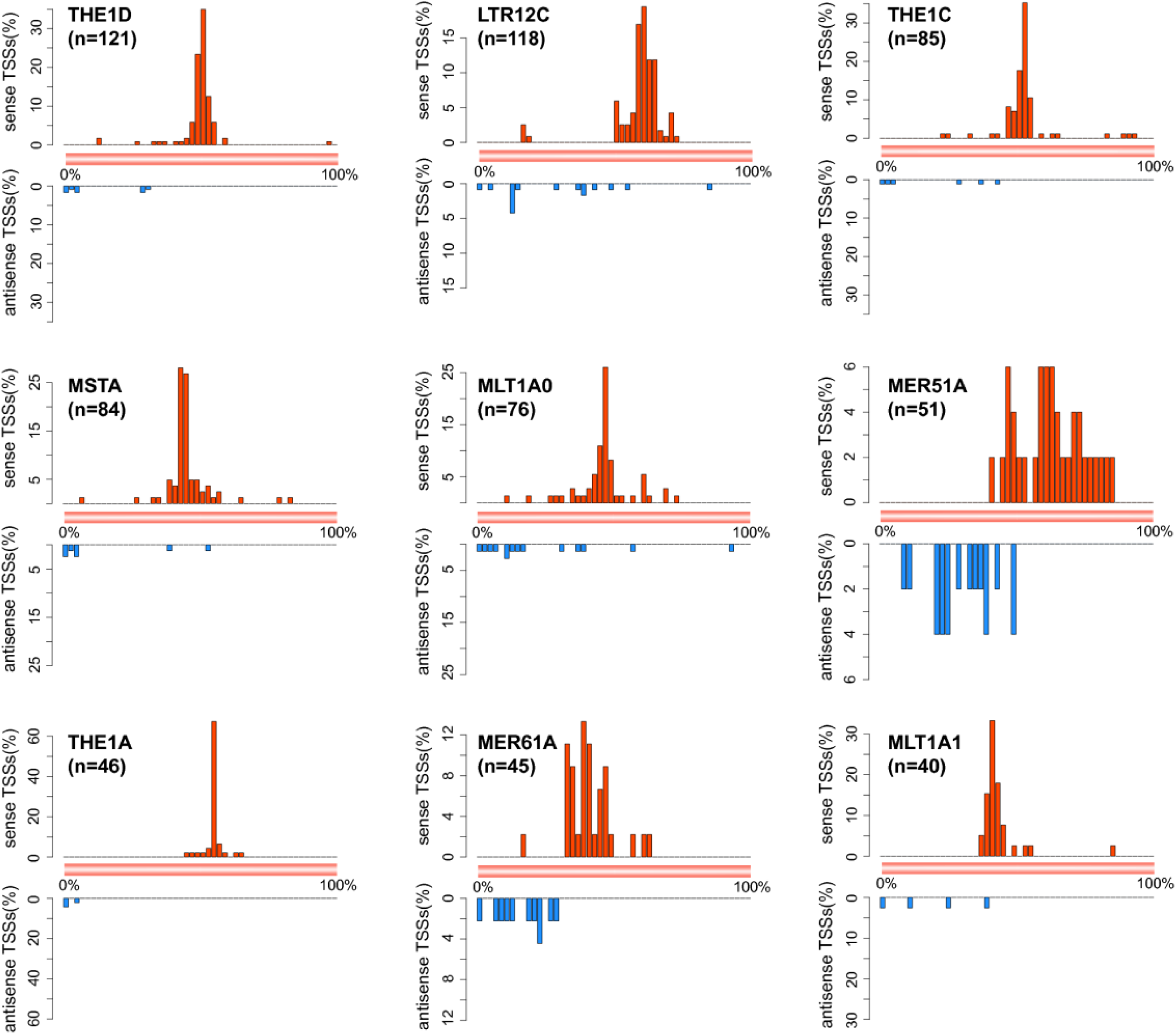
Distribution of young TSSs along LTR subfamilies. These nine subfamilies are among the top 10 LTR subfamilies which harbor most young TSSs. The tenth, THEIB, has already been shown in **Fig. 2b**. Number of young TSSs for each subfamily is given in the bracket.

**Figure S4.**
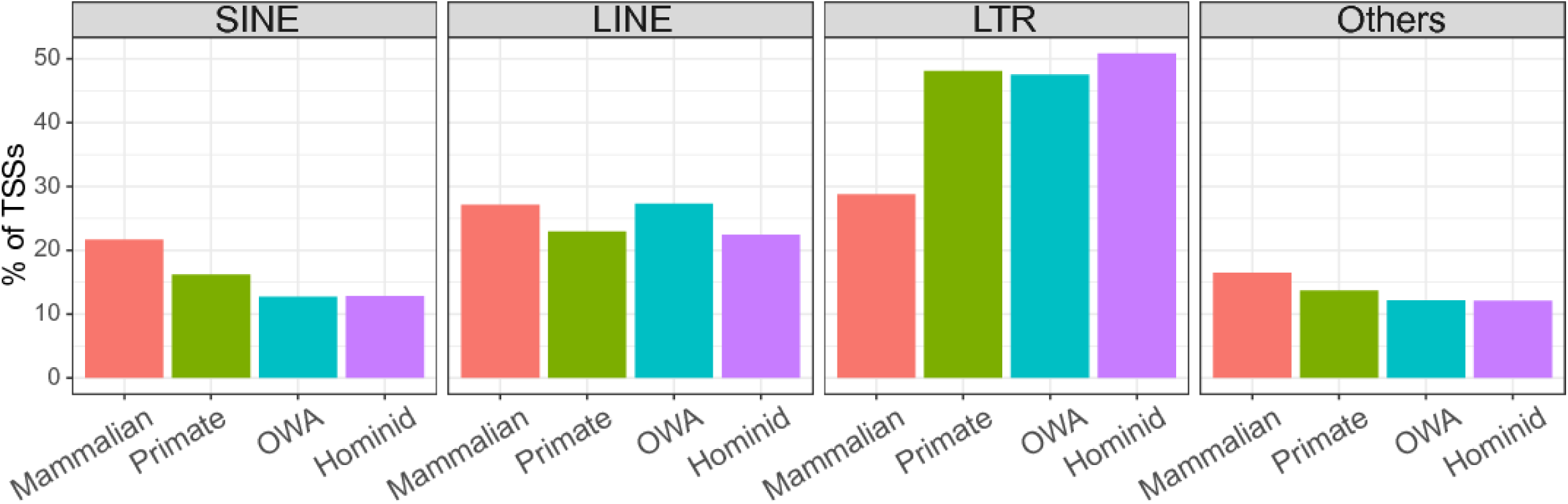
Percentages of TSSs associated with different retrotransposons which contain a TATA-box motif starting at 25-35 bp upstream regions of the dominant TSSs.

**Figure S5.**
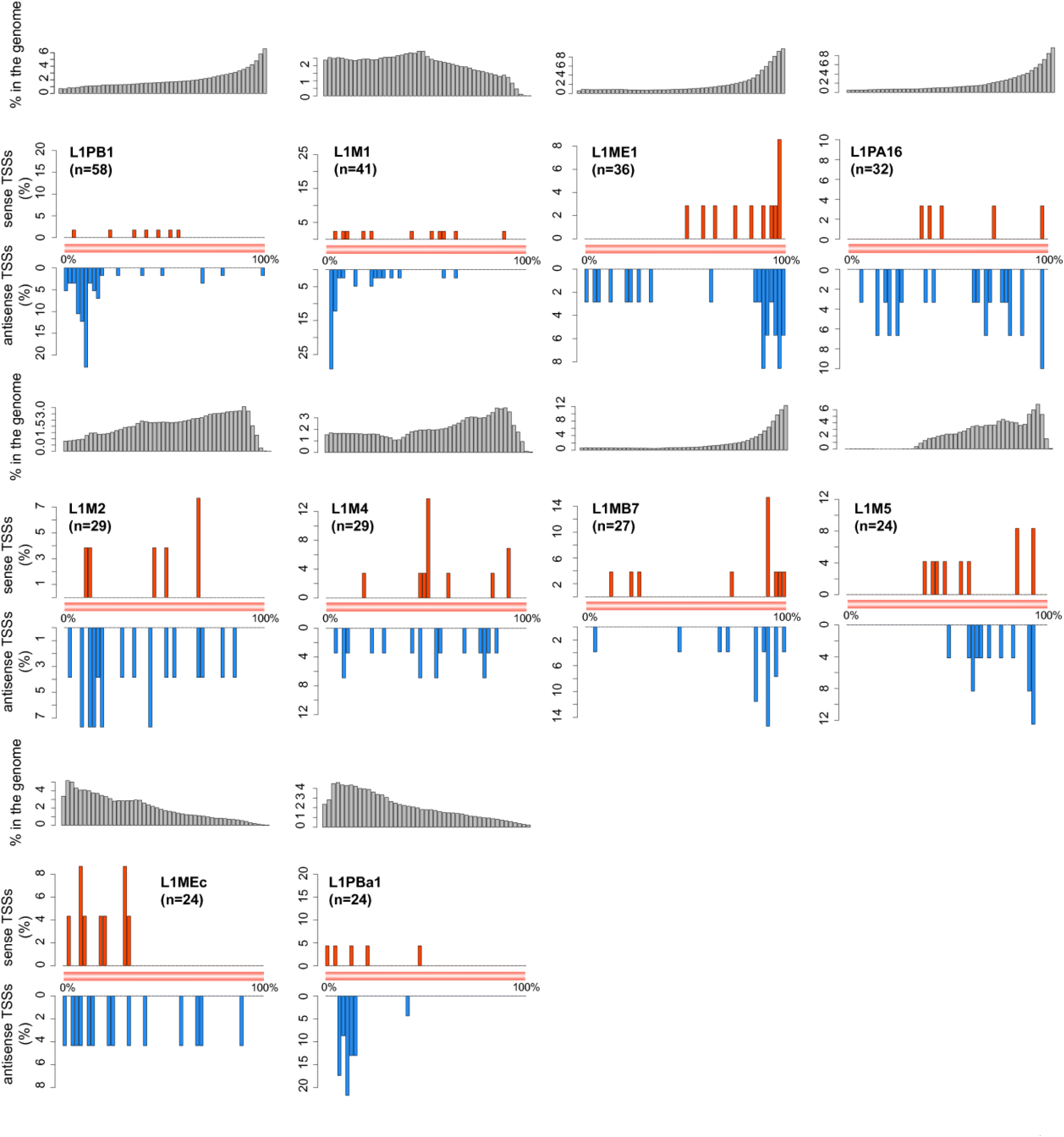
Distribution of young TSSs along L1 subfamilies. The top 10 L1 subfamilies, which harbor most young TSSs, show considerable heterogeneity regarding the positions of young TSSs within the consensus sequences. The gray barplots are background positional distributions of sequences from the corresponding subfamilies in the human genome. Number of young TSSs for each subfamily is given in the bracket.

**Figure S6.**
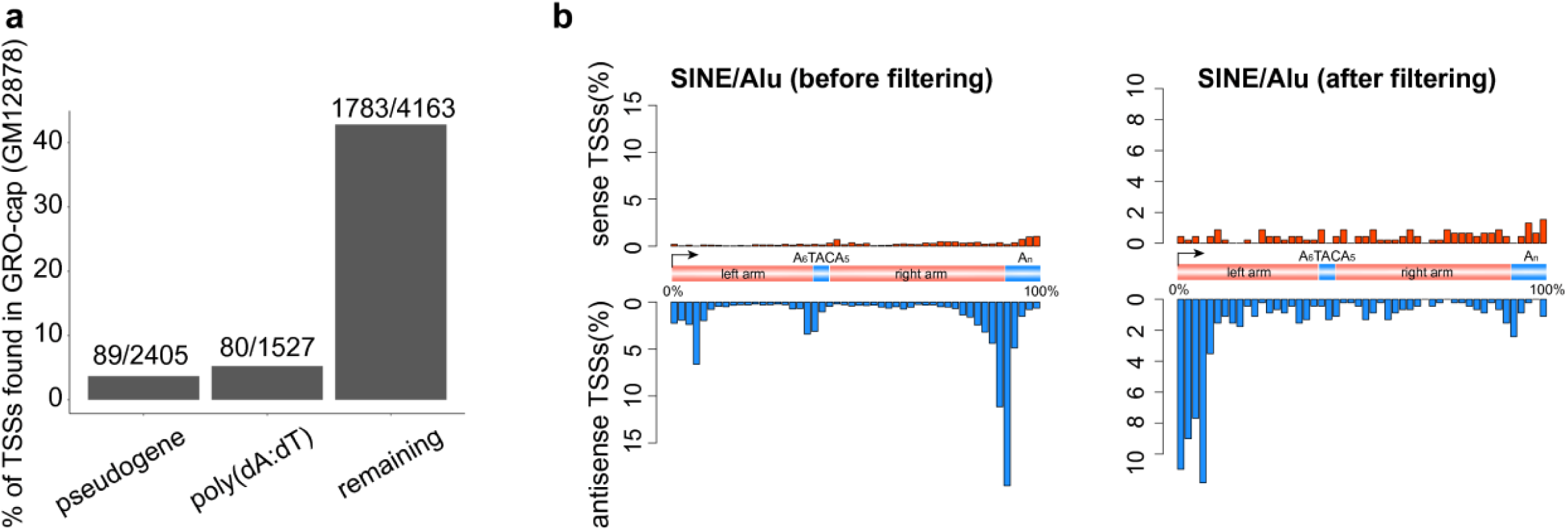
Putative false positives associated with pseudogenes and poly(dA:dT) tracts in FANTOM 5 TSSs. (**a**) Percentages of FANTOM 5 TSSs of GM12878 found in GRO-cap defined TSSs of GM12878 (from Core et al. 2014), based on the FANTOM TSSs found only in primate lineages. A FANTOM TSS is considered to be found in the GRO-cap dataset if it is within 100 bp of a GRO-cap TSS. (**b**) Distribution of FANTOM 5 TSSs along the Alu consensus element before and after filtering the suspicious TSSs.

**Figure S7.**
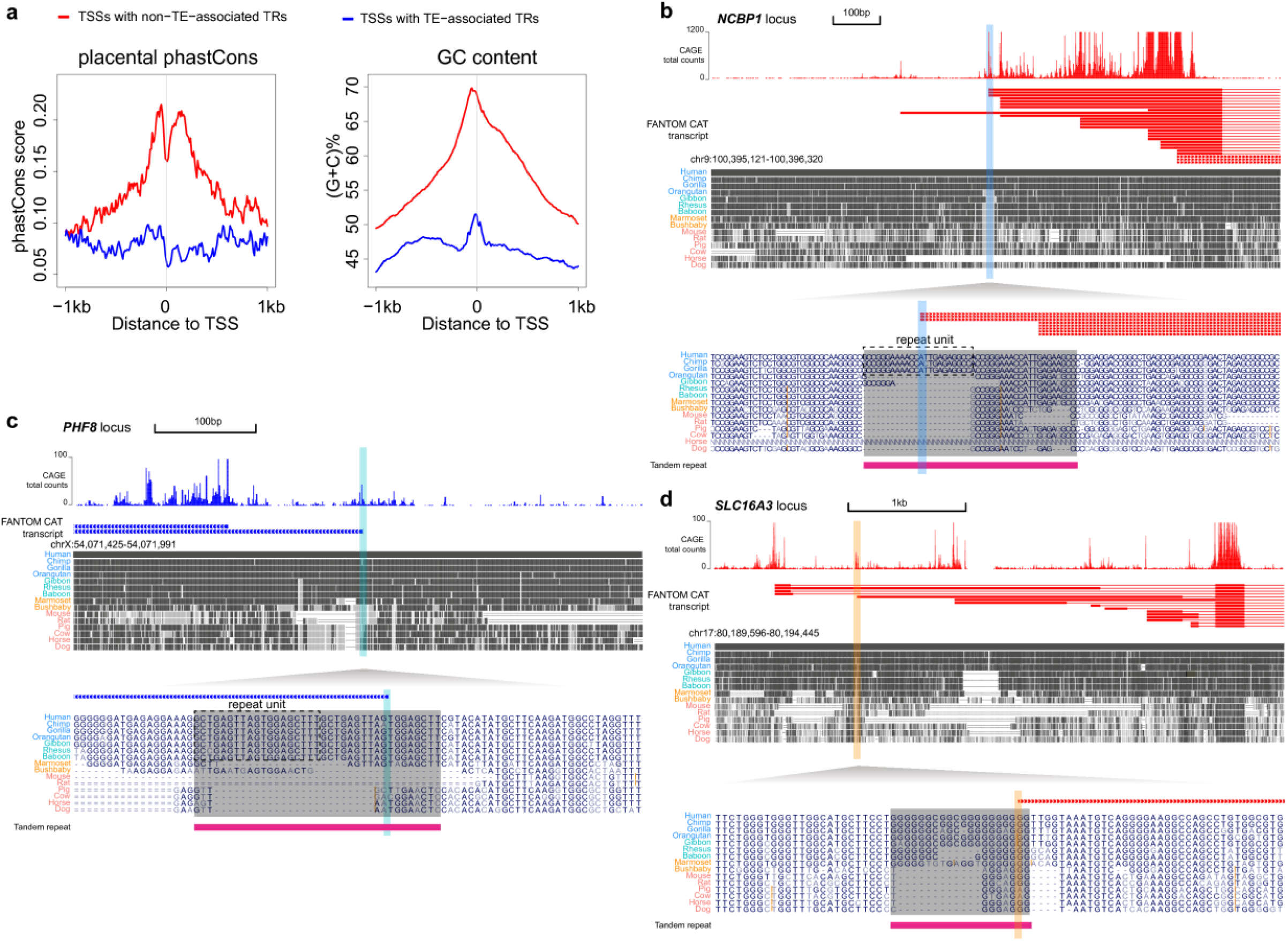
TSSs associated with tandem repeats (TRs) but not associated with TEs. (**a**) Comparison of TSSs with non-TE-associated TRs and TSSs with TE-associated TRs regarding sequence conservation scores among placental mammals and GC content. (**b-d**) Examples of non-TE-associated TR expansions which contribute to new TSSs in (b) ‘hominid’, (c) ‘OWA’ and (d) ‘primate’ groups respectively. In each panel, at the top are the CAGE total tag counts and transcripts from FANTOM (red, forward strand; blue, reverse strand) and genomic alignment blocks from UCSC genome browser; at the bottom is the enlarged region of the young TSS, with grey shade indicating the TR expansion.

**Figure S8.**
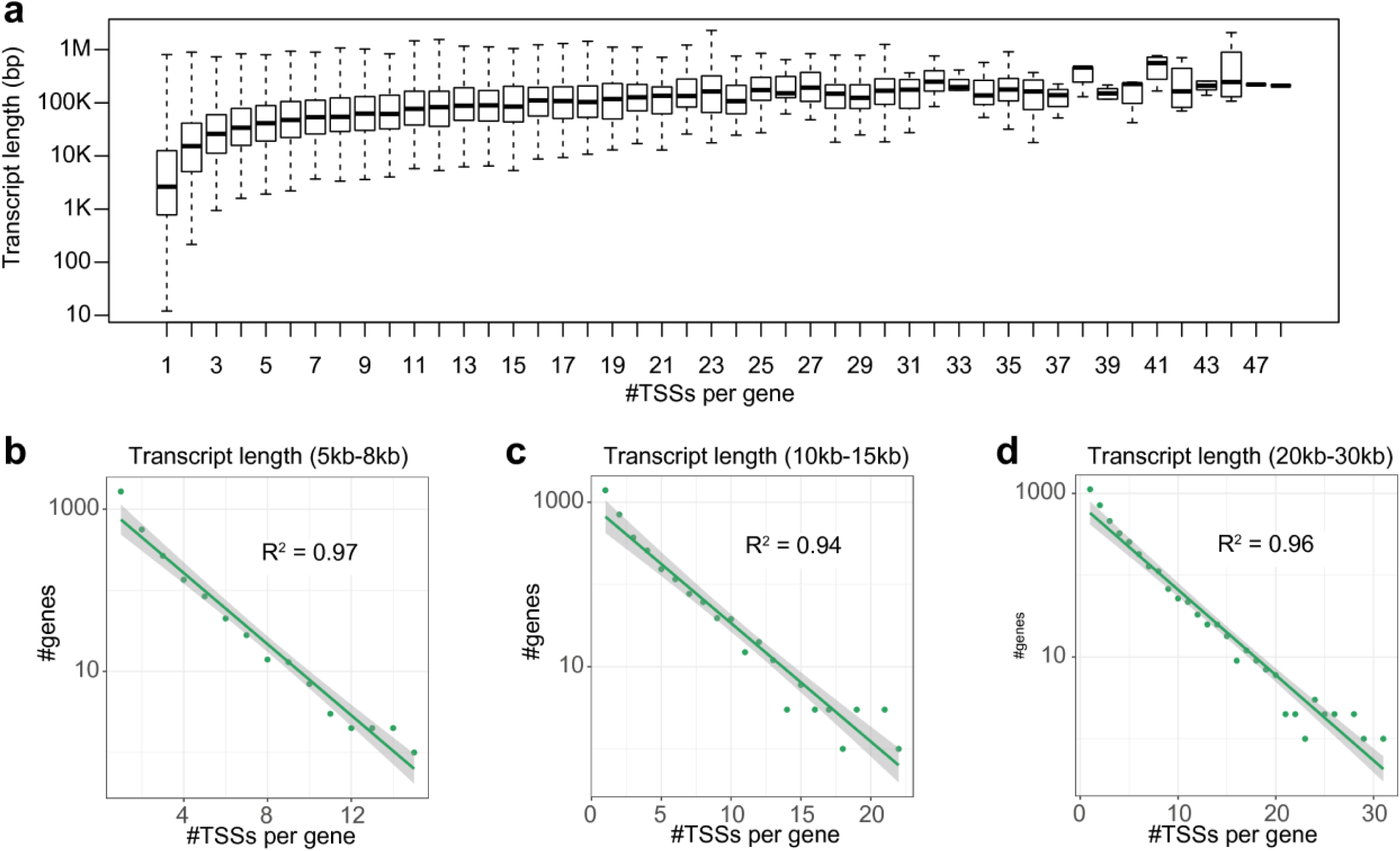
The exponential relationship between number of genes with a specific number of TSSs and number of TSSs per gene is independent of the gene lengths. (**a**) Boxplots of transcript lengths for genes with different numbers of TSSs. For each gene, we used the length of its longest transcript. Although genes that have more TSSs tend to have longer transcripts, there are also many long genes that have small numbers of TSSs. Inspecting genes within specific length ranges (panels **b-d**), still reveals a clear exponential relationship. R^2^ is the coefficient of determination for the linear regression in the figure.

**Figure S9.**
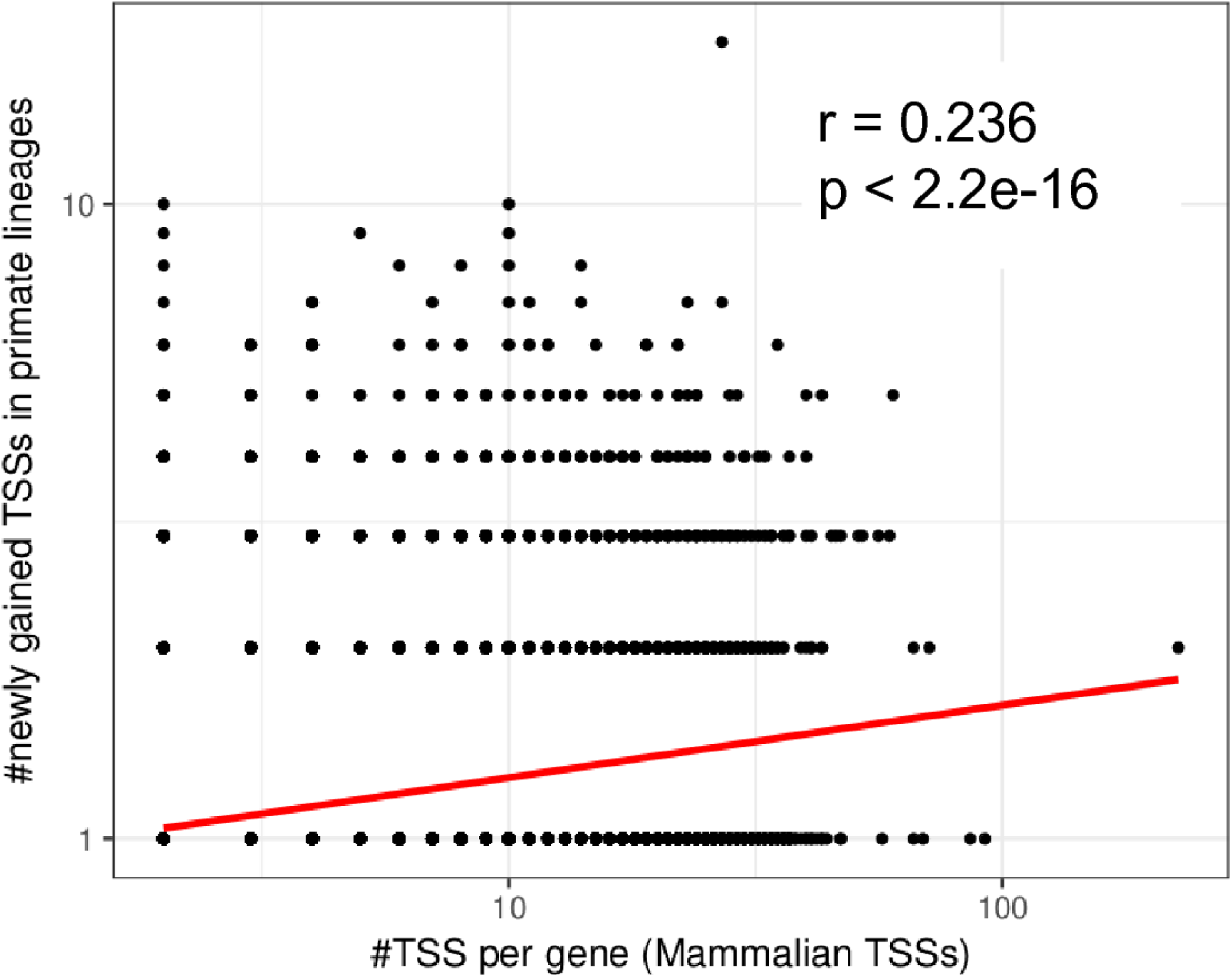
Relationship between the number of old (‘mammalian’) TSSs per gene and the number of newly gained TSSs in primate lineages, on a log10 scale. The red line is derived from linear regression based on the data points. Pearson’s r and the corresponding p-value are also shown in the figure.

**Figure S10.**
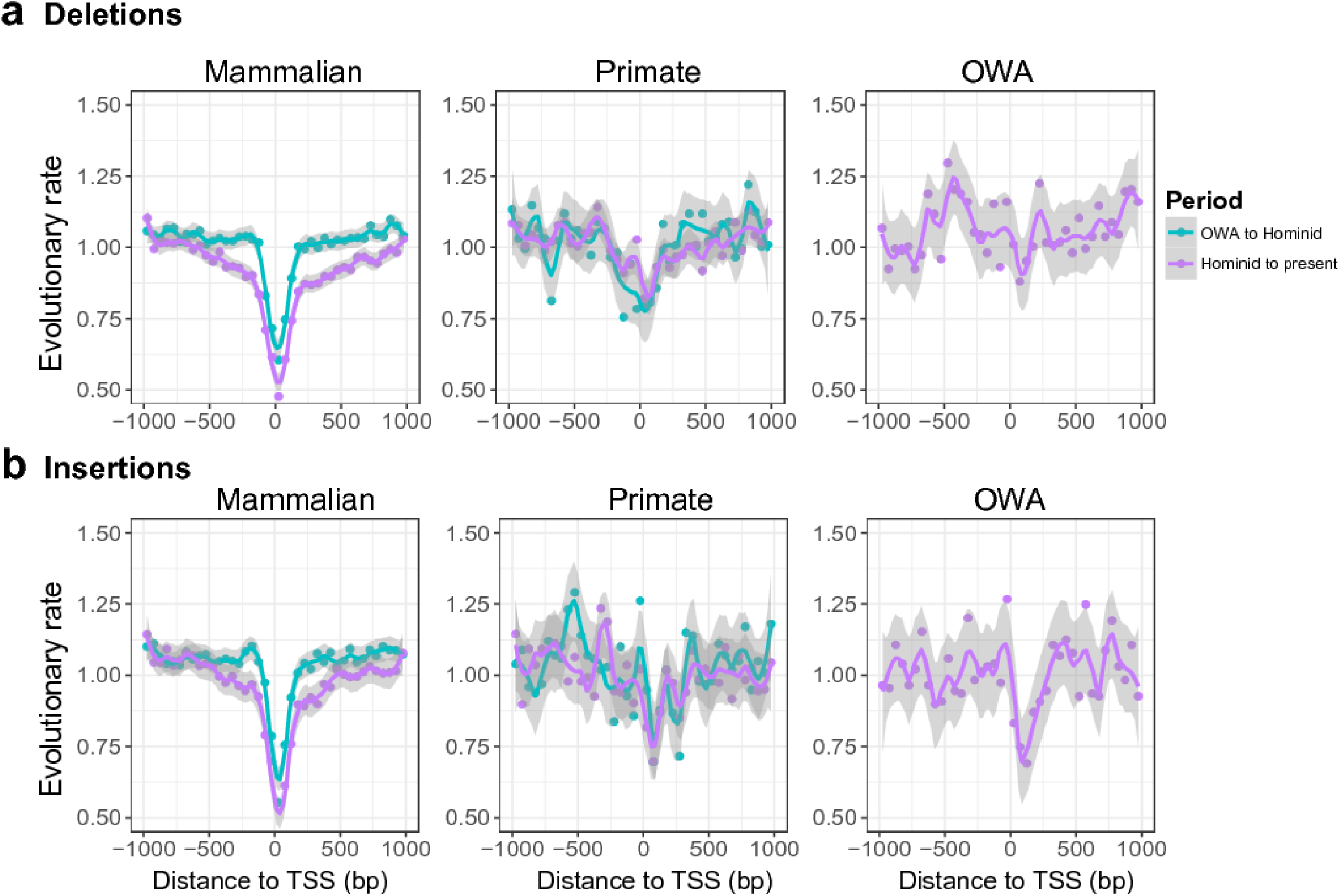
Relative deletion and insertion rates (normalized by genomic average) inferred from genomic alignments for three TSS groups. Average rate were calculated for 40 bins along TSS±1kb. We estimated insertion/deletion rates of two periods for ‘mammalian’ and ‘primate’ groups, but only one for the ‘OWA’ group so as to focus on the evolutionary rates after TSS loci emerged in the genome. Fitting curves were estimated by ‘loess’ method.

**Figure S11.**
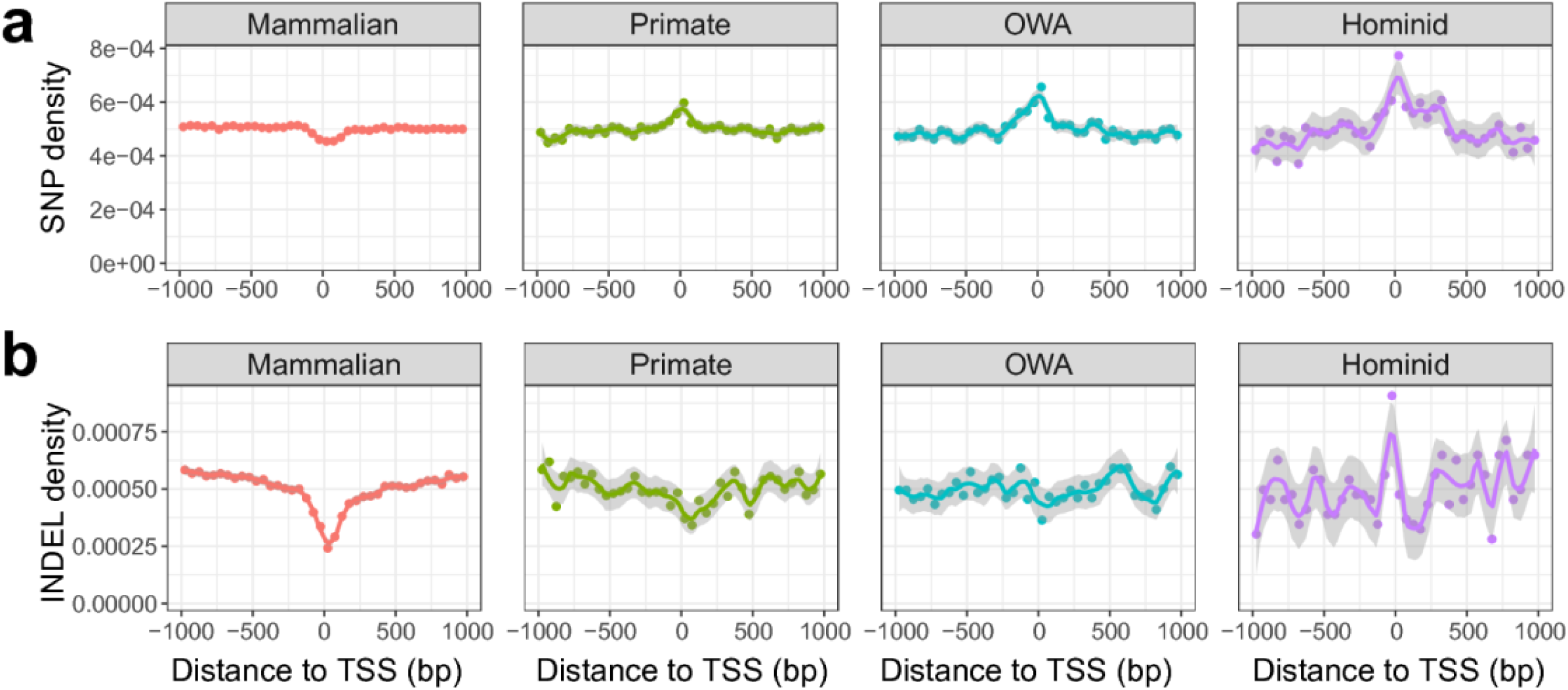
Single nucleotide polymorphism (SNP, panel a) and insertion/deletion (INDEL, panel b) densities around TSSs (40 bins along TSS±1kb), based on variants of the 1000 genomes project phase 3 release. Only the biallelic variants with a minor allele frequency of ≥ 0.01 were considered. Because the genotype files in 1000 genomes project lack the ancestral allele information for insertion/deletion variants, the insertion and deletion variants were merged together for this analysis. Fitting curves were estimated by the ‘loess’ method.

**Figure S12.**
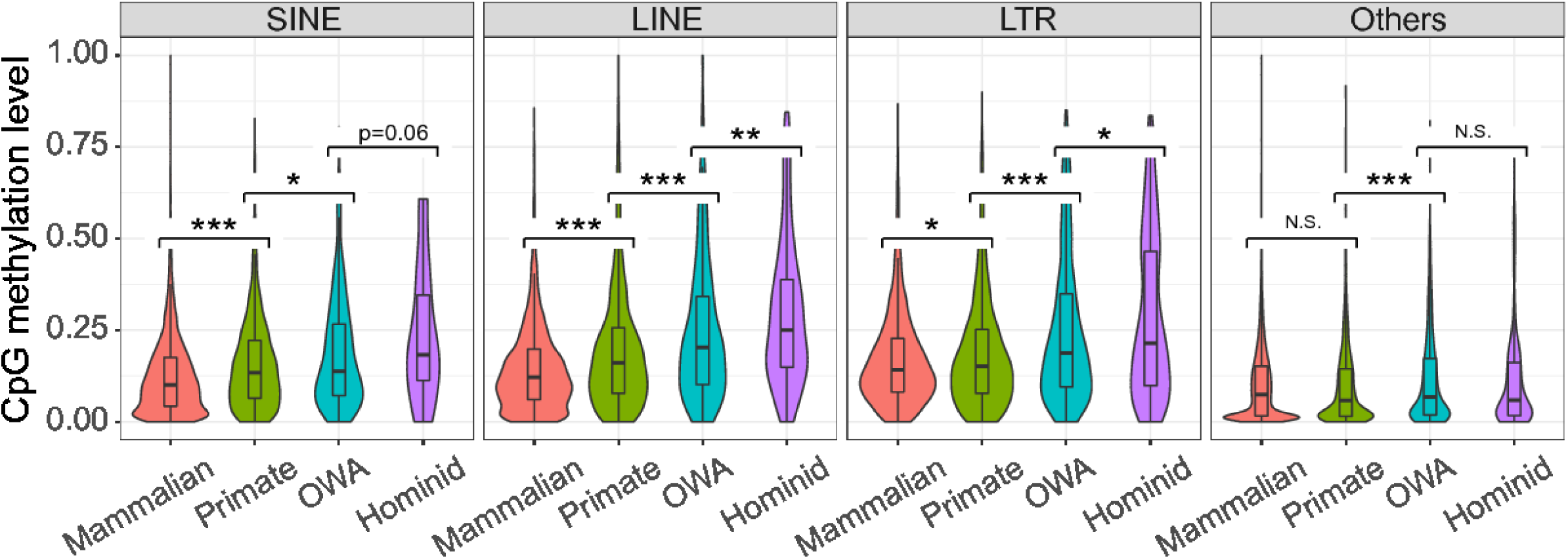
DNA methylation in TSS loci in the germline. Violin and box plots for germline CpG methylation levels (data from Guo et al. 2015) in different TSS subgroups defined by the types of associated retrotransposons. For each TSS, average methylation level of CpGs in the 2 kb around the TSS was calculated. The TSSs in the “Others” group are mostly non-TE-associated TSSs, except for a few that are associated with DNA transposons. Statistical significance was calculated using the one tailed Wilcoxon rank sum test (“*”, p < 0.05; “**”, p < 0.01; “***”, p < 0.001; N.S., not significant).

**Figure S13.**
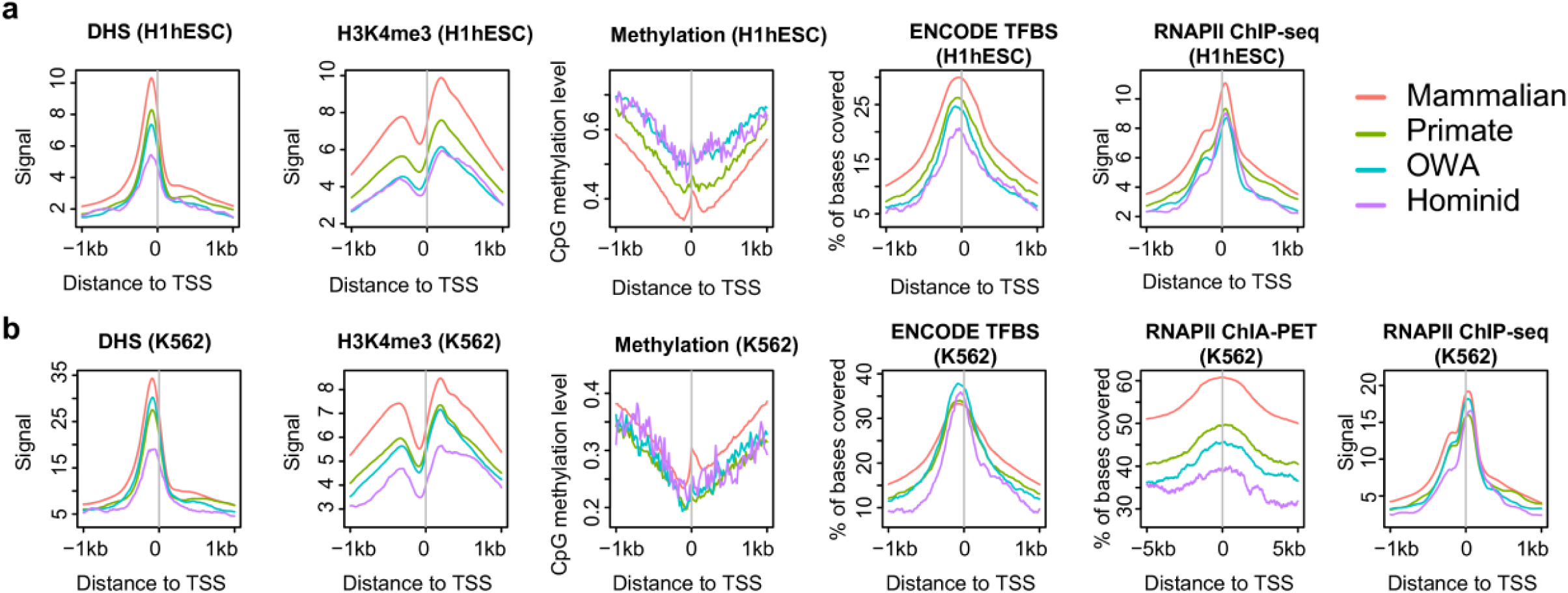
(**a**) Meta-profiles of functional signatures in H1-hESC cell line in different TSS groups. (**b**) Meta-profiles of functional signatures in K562 cell line in different TSS groups. Global hypomethylation in the K562 cell line has been previously reported, so the similar pattern of DNA methylation meta-profiles in K562 across TSS groups is not surprising. For the TFBS analysis, we merged the called peaks of TF ChIP-seq datasets and calculated how many bases around TSSs are covered by the peaks. The figure for RNAP II ChIA-PET in H1-hESC is missing because of lack of publicly available data.

**Figure S14.**
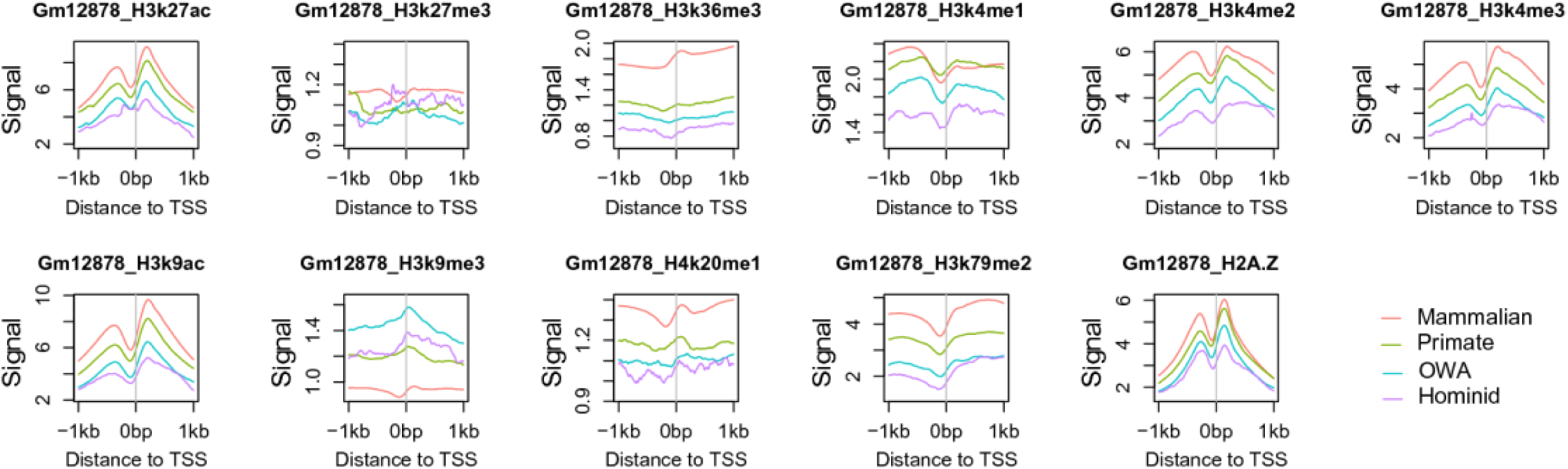
Meta-profiles for histone modifications in GM12878, supplementary to that shown in Fig. 4. All the data was obtained from ENCODE project.

**Figure S15.**
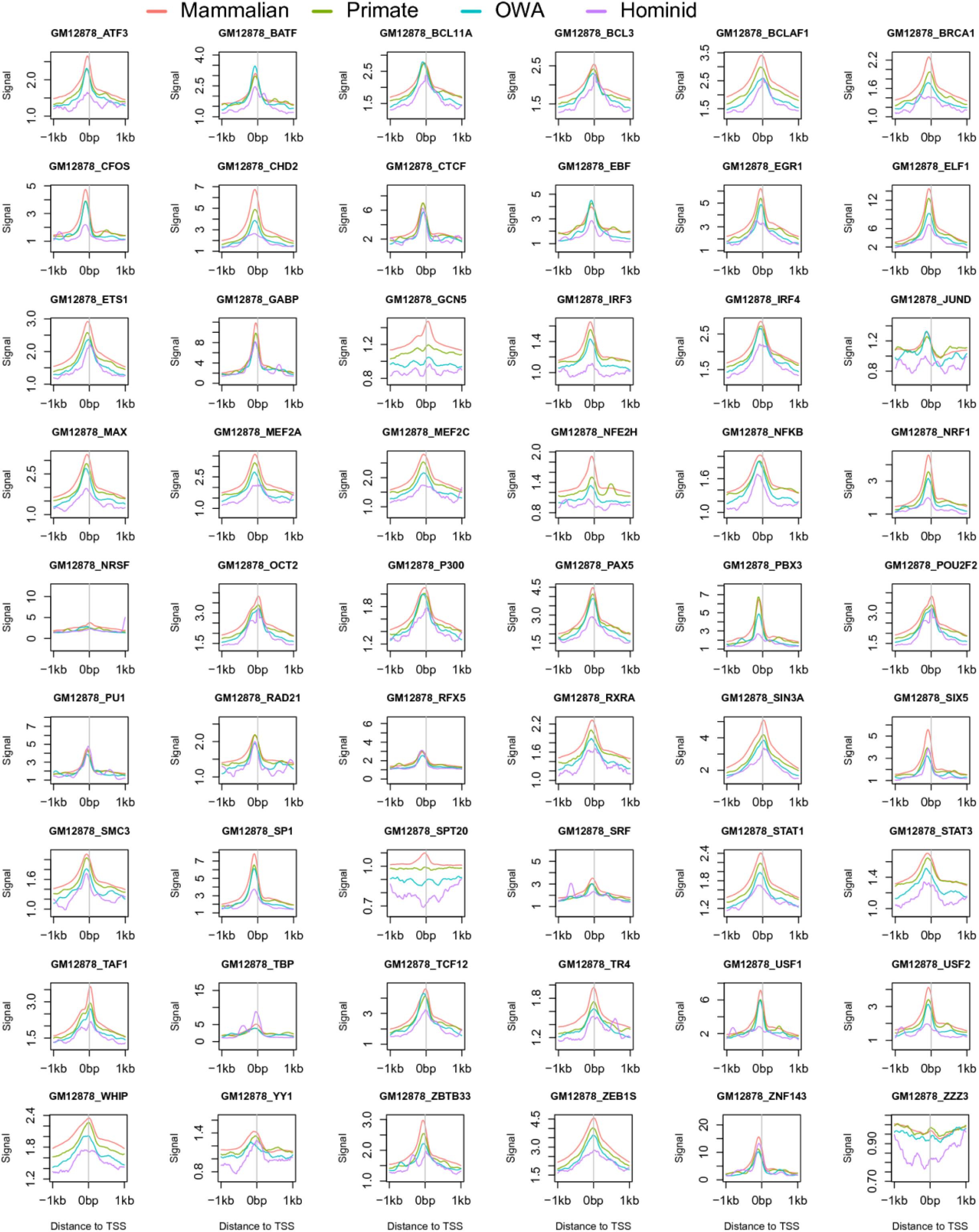
Meta-profiles for TF ChIP-seq signals in GM12878 cell line in different TSS groups. All the data was obtained from ENCODE project.

**Figure S16.**
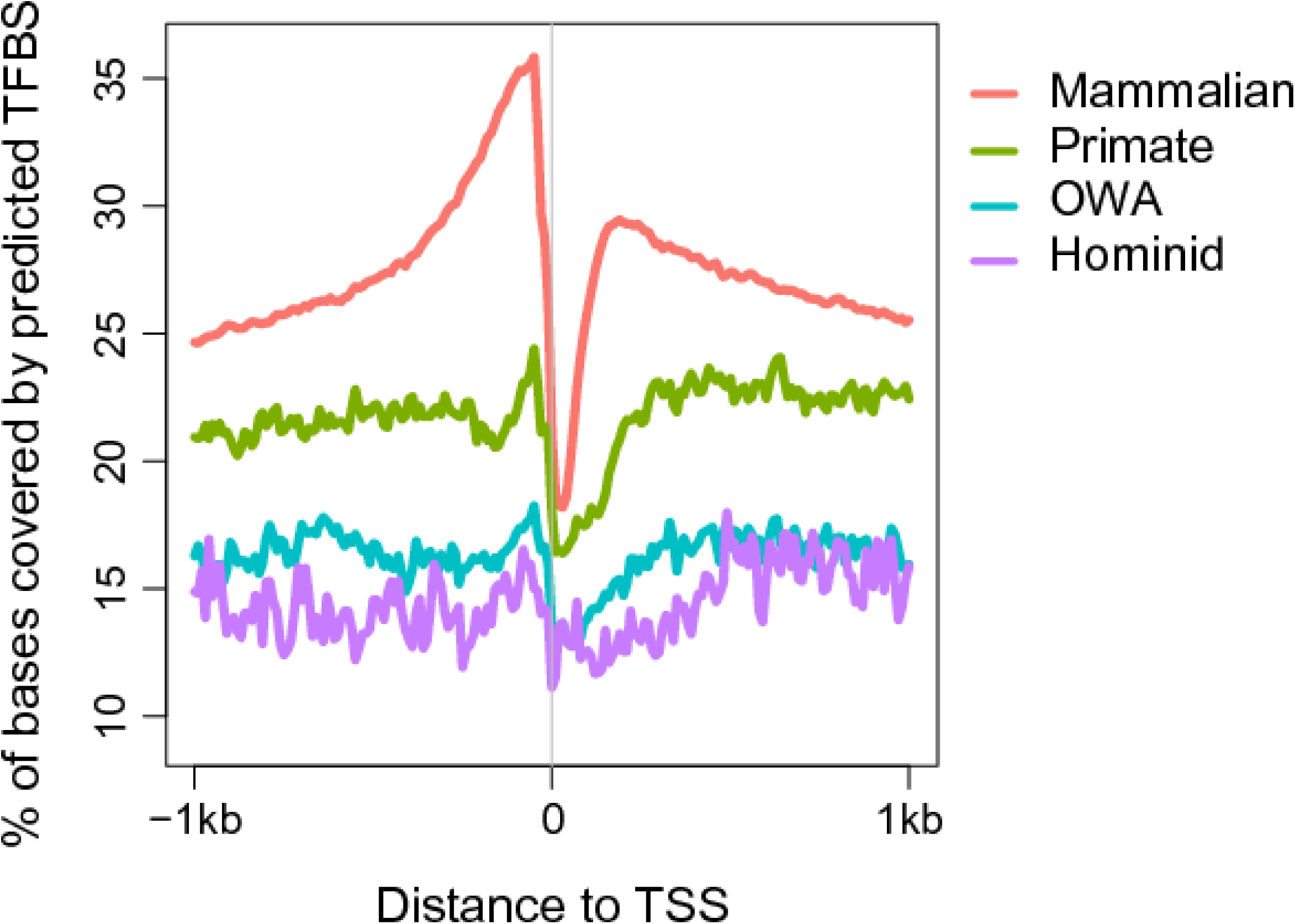
Comparison of the coverage by computationally predicted TFBSs between four TSS groups. The computationally predicted TFBSs in human genome were from ENCODE project (http://compbio.mit.edu/encode-motifs/). Note that the TFBSs predicted by computational methods are based on binding motifs, usually smaller than the called peaks in the TF ChIP-seq data that were used for generating **Fig. 4**.

**Figure S17.**
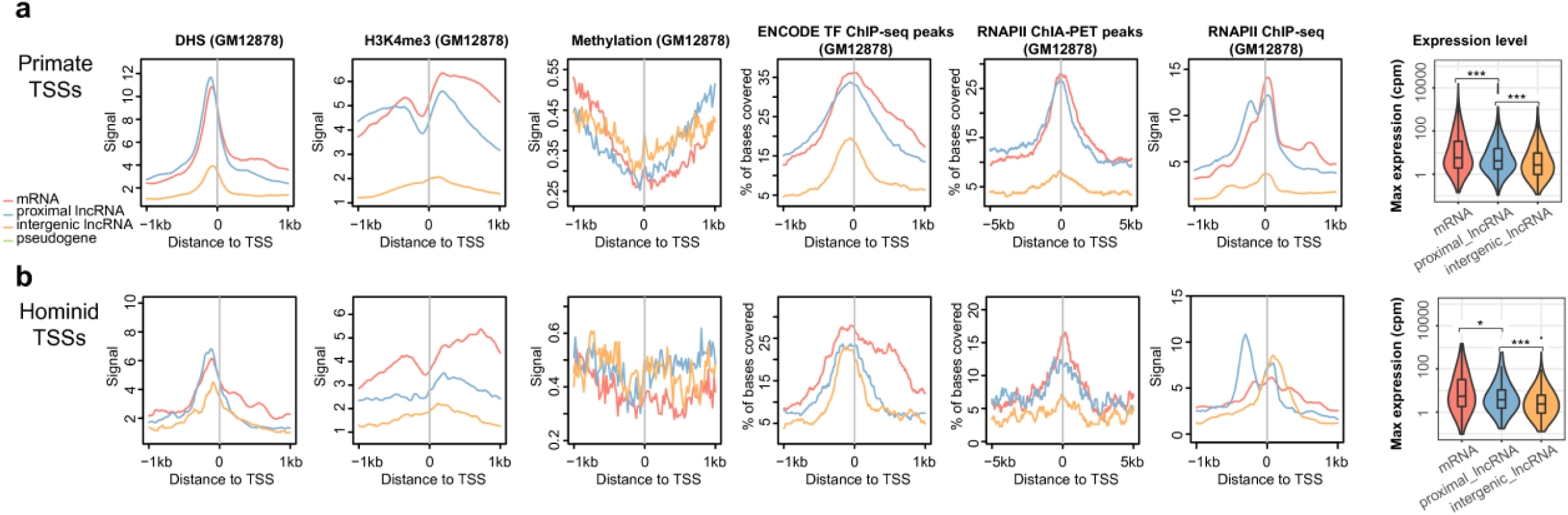
Meta-profiles of functional signatures in GM12878 cell line for different TSS subgroups, defined by transcript types. (**a**) For ‘primate’ TSS subgroups. (**b**) For ‘hominid’ TSS subgroups. Statistical significance was calculated using the one-tailed Wilcoxon rank sum tests (“*”, p < 0.05; “**”, p < 0.01; “***”, p < 0.001).

**Figure S18.**
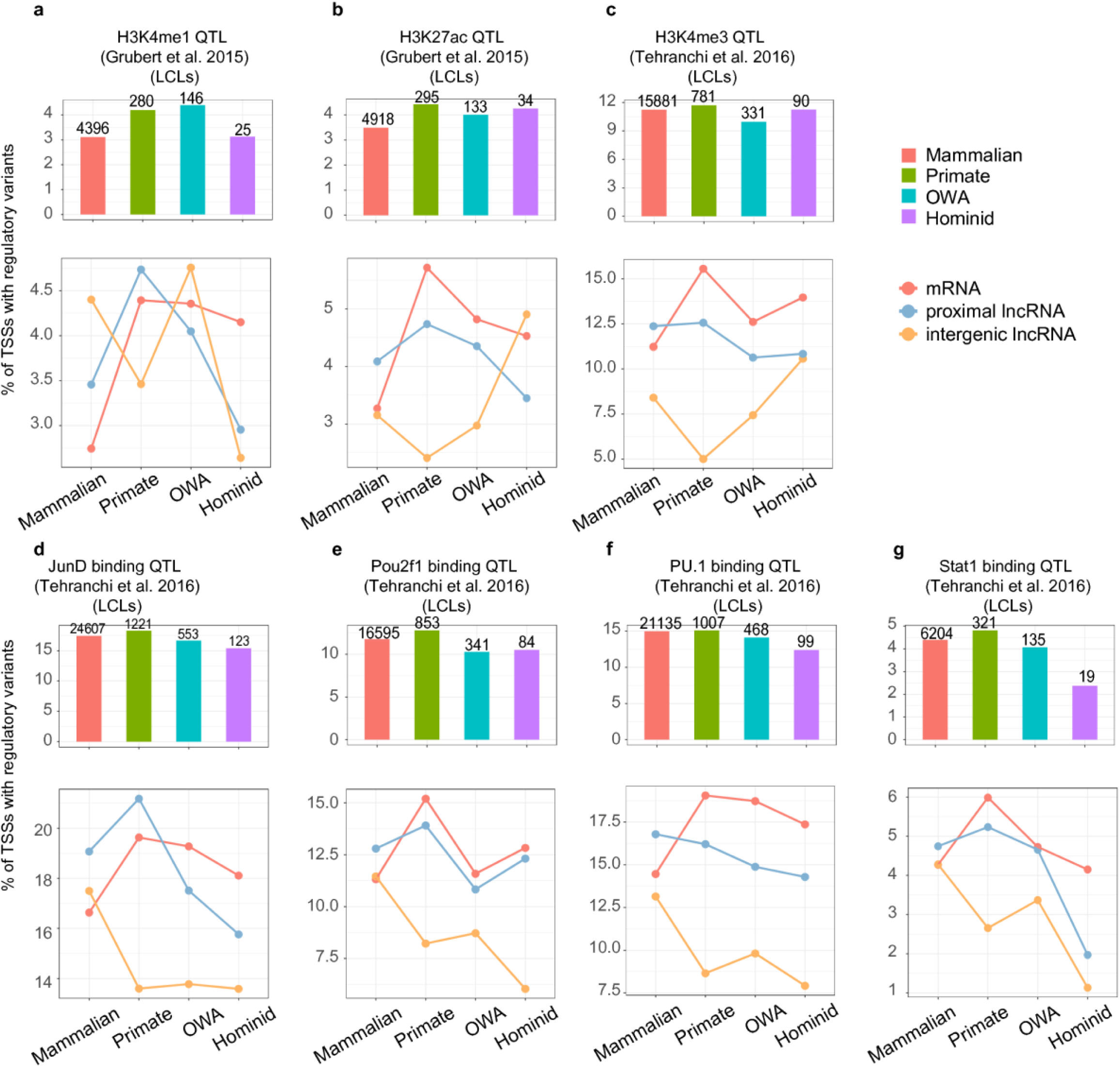
Proportions of TSSs harboring regulatory variants within TSS±1kb in different TSS groups in additional datasets. The results in this figure were based on regulatory variants with derived allele frequency (DAF) ≥ 0.01. Above the bars are the numbers of TSSs with regulatory variants. Note that for the H3K4me3 QTL dataset from Grubert et al. (2015), the numbers of regulatory variants found in the TSS groups/subgroups are very small, so the trends shown in the panels **a-b** for different transcript types may not accurately reflect actual trends. LCLs, lymphoblastoid cell lines.

**Figure S19.**
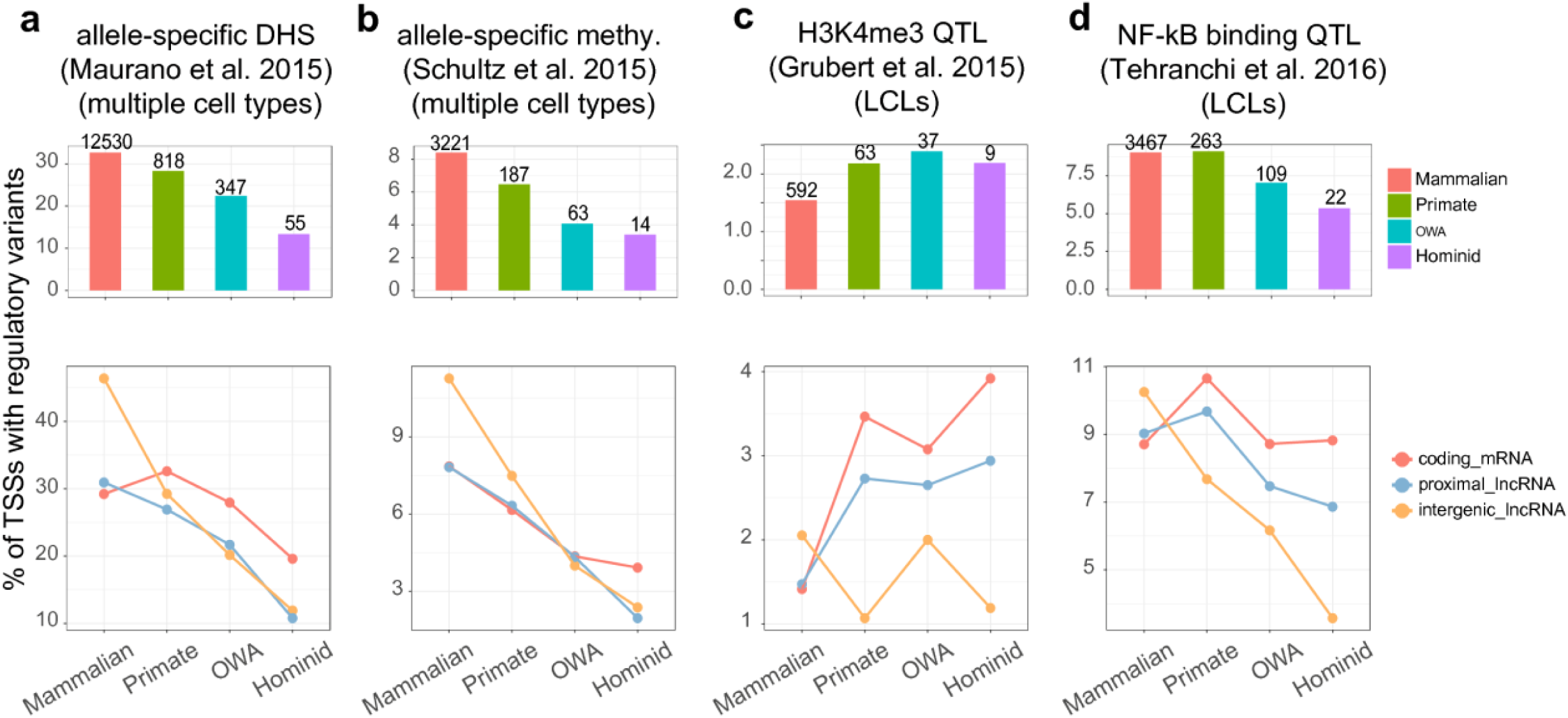
Proportions of TSSs harboring regulatory variants within TSS±1kb in different TSS groups excluding the TSSs separated by < 2 kb. The shown results are based on variants with DAF ≥ 0.01. Above the bars are the numbers of TSSs with regulatory variants. Note that for the H3K4me3 QTL dataset from Grubert et al. (2015), the numbers of regulatory variants found in the TSS groups/subgroups are very small, so the changing trends shown in the panel **c** for different transcript types may not accurately reflect actual trends. LCLs, lymphoblastoid cell lines.

**Figure S20.**
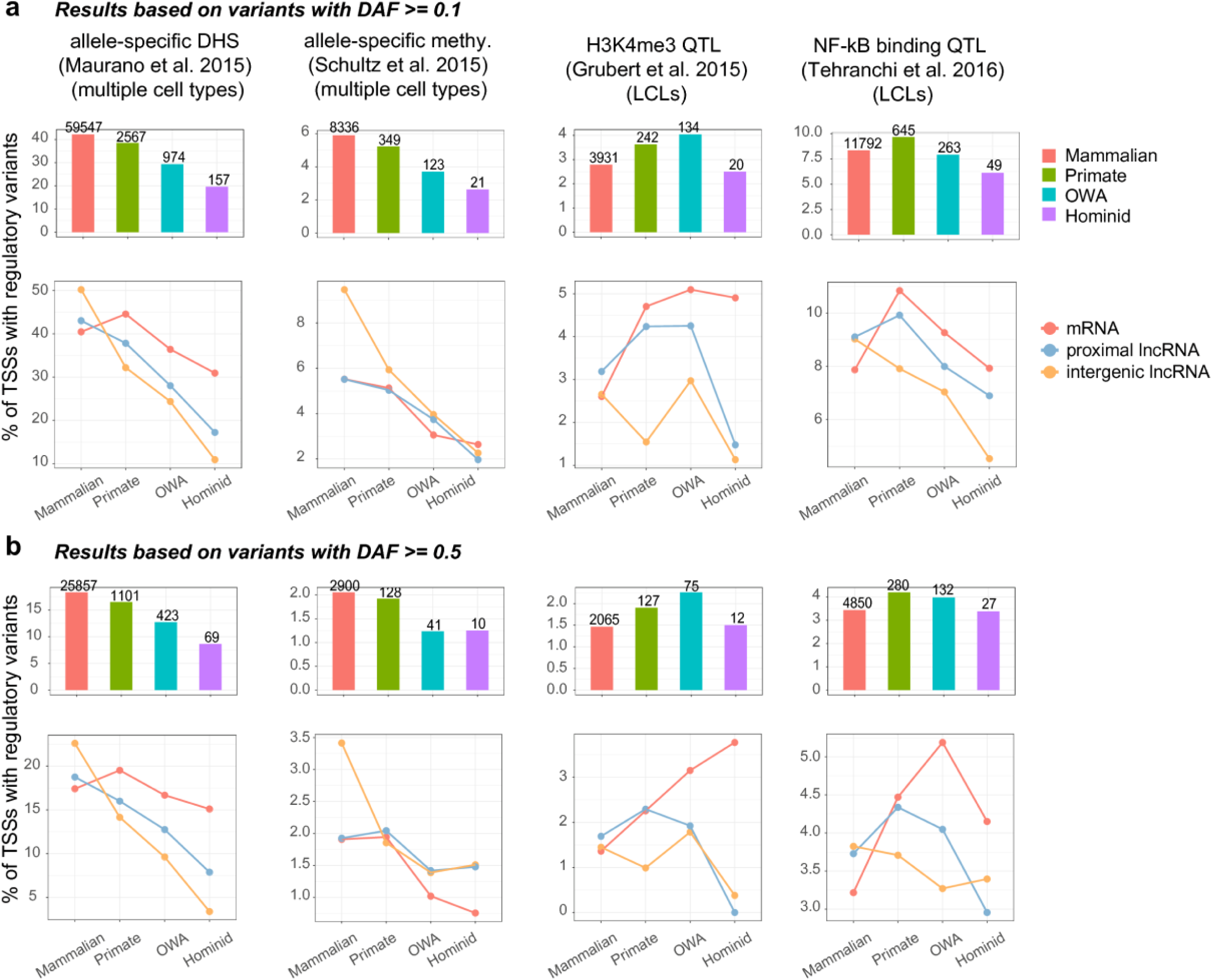
Proportions of TSSs harboring regulatory variants within TSS±1kb in different TSS groups, based on variants with higher thresholds of derived allele frequency (DAF). (**a**) Results based on variants with DAF ≥ 0.1. (**b**) Results based on variants with DAF ≥ 0.5. Above the bars are the numbers of TSSs with regulatory variants. Note that for the results based on variants with DAF ≥ 0.5, the numbers of regulatory variants found in the TSS groups/subgroups are very small, so the changing trends shown in the some panels for different transcript types may not accurately reflect actual trends. LCLs, lymphoblastoid cell lines.

**Figure S21.**
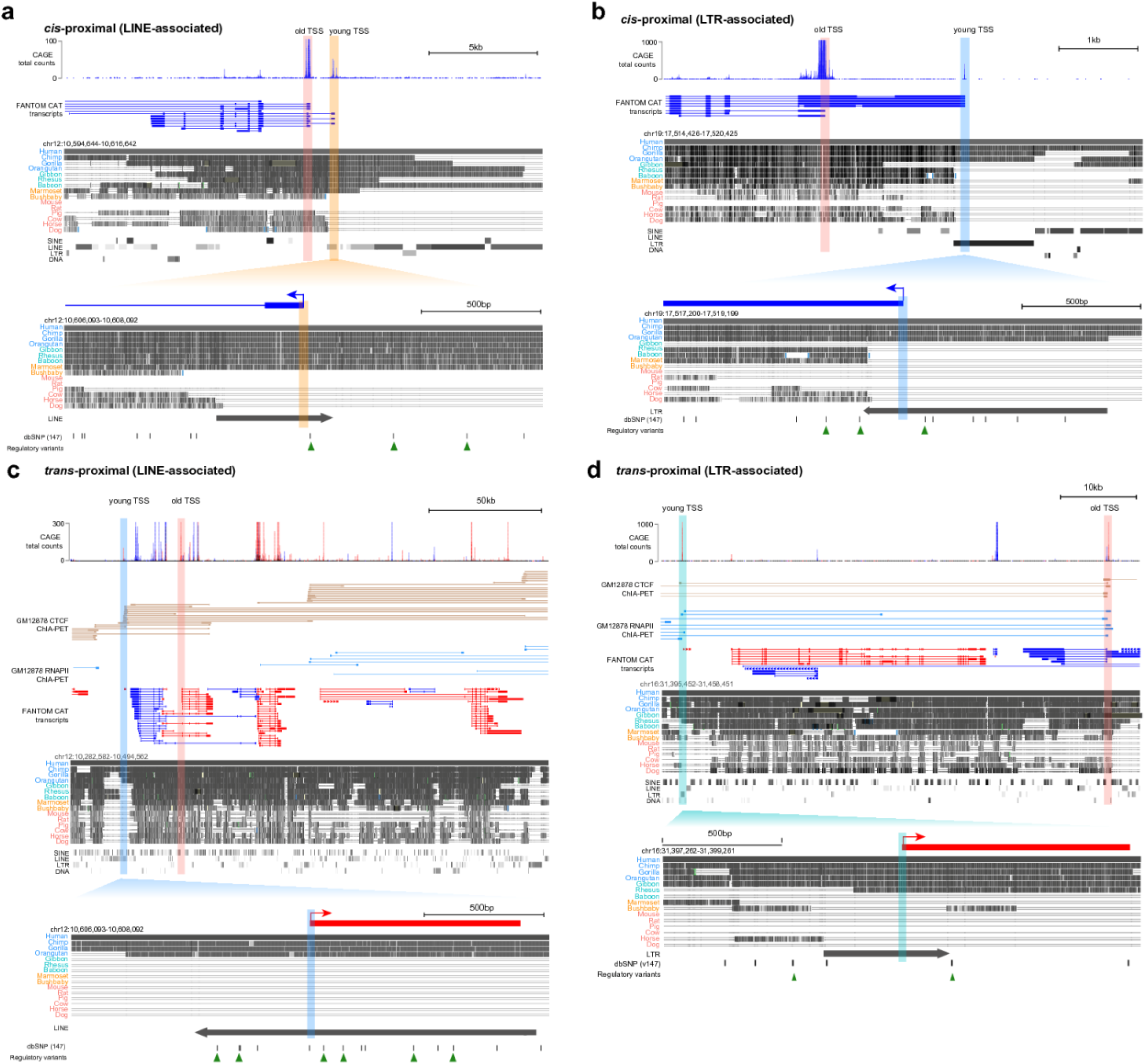
Additional examples for *cis*-proximal and *trans*-proximal young TSSs. In each panel, from top to bottom: 1) CAGE total tag counts from FANTOM; 2) CTCF and RNAP II ChIA-PET interactions (only for *trans*-proximal examples); 3) FANTOM CAT transcript models, red for forward-strand and blue for reverse-strand transcripts; 4) genome alignments represented by grey blocks and transposable elements within this region, generated from UCSC genome browser 5) the enlarged region of the young TSS. The old and young TSSs are indicated with shades of different colors (red, “mammalian”; green, “primate”; cyan, “OWA”; blue, “hominid”). The positions of regulatory variants are shown with small triangles in the enlarged figures.

## References

1 Clark, M. B. et al. The reality of pervasive transcription. PLoS biology 9, e1000625; discussion e1001102, doi:10.1371/journal.pbio.1000625 (2011).s

2 Wade, J. T. & Grainger, D. C. Pervasive transcription: illuminating the dark matter of bacterial transcriptomes. Nature reviews. Microbiology 12, 647–653, doi:10.1038/nrmicro3316 (2014).

3 Djebali, S. et al. Landscape of transcription in human cells. Nature 489, 101– 108, doi:10.1038/nature11233 (2012).

4 FANTOM Consortium et al. A promoter-level mammalian expression atlas. Nature 507, 462–470, doi:10.1038/nature13182 (2014).

5 Core, L. J. et al. Analysis of nascent RNA identifies a unified architecture of initiation regions at mammalian promoters and enhancers. Nature genetics 46, 1311–1320, doi:10.1038/ng.3142 (2014).

6 Core, L. J., Waterfall, J. J. & Lis, J. T. Nascent RNA sequencing reveals widespread pausing and divergent initiation at human promoters. Science 322, 1845–1848, doi:10.1126/science.1162228 (2008).

7 Kim, T. K. & Shiekhattar, R. Architectural and Functional Commonalities between Enhancers and Promoters. Cell 162, 948–959, doi:10.1016/j.cell.2015.08.008 (2015).

8 Li, W., Notani, D. & Rosenfeld, M. G. Enhancers as non-coding RNA transcription units: recent insights and future perspectives. Nature reviews. Genetics 17, 207–223, doi:10.1038/nrg.2016.4 (2016).

9 Andersson, R. et al. An atlas of active enhancers across human cell types and tissues. Nature 507, 455–461, doi:10.1038/nature12787 (2014).

10 Frith, M. C. et al. Evolutionary turnover of mammalian transcription start sites. Genome research 16, 713–722, doi:10.1101/gr.5031006 (2006).

11 Main, B. J., Smith, A. D., Jang, H. & Nuzhdin, S. V. Transcription start site evolution in Drosophila. Molecular biology and evolution 30, 1966–1974, doi:10.1093/molbev/mst085 (2013).

12 Yokoyama, K. D., Thorne, J. L. & Wray, G. A. Coordinated genome-wide modifications within proximal promoter cis-regulatory elements during vertebrate evolution. Genome biology and evolution 3, 66–74, doi:10.1093/gbe/evq078 (2011).

13 Young, R. S. et al. The frequent evolutionary birth and death of functional promoters in mouse and human. Genome research 25, 1546–1557, doi:10.1101/gr.190546.115 (2015).

14 Taylor, M. S. et al. Heterotachy in mammalian promoter evolution. PLoS genetics 2, e30, doi:10.1371/journal.pgen.0020030 (2006).

15 Ward, L. D. & Kellis, M. Evidence of abundant purifying selection in humans for recently acquired regulatory functions. Science 337, 1675–1678, doi:10.1126/science.1225057 (2012).

16 Scala, G., Affinito, O., Miele, G., Monticelli, A. & Cocozza, S. Evidence for evolutionary and nonevolutionary forces shaping the distribution of human genetic variants near transcription start sites. PloS one 9, e114432, doi:10.1371/journal.pone.0114432 (2014).

17 Schor, I. E. et al. Promoter shape varies across populations and affects promoter evolution and expression noise. Nature genetics 49, 550–558, doi:10.1038/ng.3791 (2017).

18 Faulkner, G. J. et al. The regulated retrotransposon transcriptome of mammalian cells. Nature genetics 41, 563–571, doi:10.1038/ng.368 (2009).

19 Nguyen, T. A. et al. High-throughput functional comparison of promoter and enhancer activities. Genome research 26, 1023–1033, doi:10.1101/gr.204834.116 (2016).

20 van Arensbergen, J. et al. Genome-wide mapping of autonomous promoter activity in human cells. Nature biotechnology 35, 145–153, doi:10.1038/nbt.3754 (2017).

21 Lenhard, B., Sandelin, A. & Carninci, P. Metazoan promoters: emerging characteristics and insights into transcriptional regulation. Nature reviews. Genetics 13, 233–245, doi:10.1038/nrg3163 (2012).

22 Sawaya, S. et al. Microsatellite tandem repeats are abundant in human promoters and are associated with regulatory elements. PloS one 8, e54710, doi:10.1371/journal.pone.0054710 (2013).

23 Bilgin Sonay, T. et al. Tandem repeat variation in human and great ape populations and its impact on gene expression divergence. Genome research 25, 1591–1599, doi:10.1101/gr.190868.115 (2015).

24 Tarailo-Graovac, M. & Chen, N. Using RepeatMasker to identify repetitive elements in genomic sequences. Current protocols in bioinformatics Chapter 4, Unit 4 10, doi:10.1002/0471250953.bi0410s25 (2009).

25 Benson, G. Tandem repeats finder: a program to analyze DNA sequences. Nucleic acids research 27, 573–580 (1999).

26 Willems, T. et al. The landscape of human STR variation. Genome research 24, 1894–1904, doi:10.1101/gr.177774.114 (2014).

27 Kanamori-Katayama, M. et al. Unamplified cap analysis of gene expression on a single-molecule sequencer. Genome research 21, 1150–1159, doi:10.1101/gr.115469.110 (2011).

28 Hon, C. C. et al. An atlas of human long non-coding RNAs with accurate 5’ ends. Nature 543, 199–204, doi:10.1038/nature21374 (2017).

29 Ohadi, M. et al. Core promoter short tandem repeats as evolutionary switch codes for primate speciation. American journal of primatology 77, 34–43, doi:10.1002/ajp.22308 (2015).

30 Gymrek, M. et al. Abundant contribution of short tandem repeats to gene expression variation in humans. Nature genetics 48, 22–29, doi:10.1038/ng.3461 (2016).

31 Slotkin, R. K. & Martienssen, R. Transposable elements and the epigenetic regulation of the genome. Nature reviews. Genetics 8, 272–285, doi:10.1038/nrg2072 (2007).

32 Merkenschlager, M. & Odom, D. T. CTCF and cohesin: linking gene regulatory elements with their targets. Cell 152, 1285–1297, doi:10.1016/j.cell.2013.02.029 (2013).

33 Pratto, F. et al. Recombination initiation maps of individual human genomes. Science 346, 1256442, doi:10.1126/science.1256442 (2014).

34 Chuong, E. B., Elde, N. C. & Feschotte, C. Regulatory activities of transposable elements: from conflicts to benefits. Nature reviews. Genetics 18, 71–86, doi:10.1038/nrg.2016.139 (2017).

35 Baudat, F., Imai, Y. & de Massy, B. Meiotic recombination in mammals: localization and regulation. Nature reviews. Genetics 14, 794–806, doi:10.1038/nrg3573 (2013).

36 Eckert, K. A. & Hile, S. E. Every microsatellite is different: Intrinsic DNA features dictate mutagenesis of common microsatellites present in the human genome. Molecular carcinogenesis 48, 379–388, doi:10.1002/mc.20499 (2009).

37 Kelkar, Y. D., Tyekucheva, S., Chiaromonte, F. & Makova, K. D. The genome-wide determinants of human and chimpanzee microsatellite evolution. Genome research 18, 30–38, doi:10.1101/gr.7113408 (2008).

38 Tang, Z. et al. CTCF-Mediated Human 3D Genome Architecture Reveals Chromatin Topology for Transcription. Cell 163, 1611–1627, doi:10.1016/j.cell.2015.11.024 (2015).

39 Albert, F. W. & Kruglyak, L. The role of regulatory variation in complex traits and disease. Nature reviews. Genetics 16, 197–212, doi:10.1038/nrg3891 (2015).

40 Cheng, C. et al. Understanding transcriptional regulation by integrative analysis of transcription factor binding data. Genome research 22, 1658–1667, doi:10.1101/gr.136838.111 (2012).

41 GTEx Consortium. Human genomics. The Genotype-Tissue Expression (GTEx) pilot analysis: multitissue gene regulation in humans. Science 348, 648–660, doi:10.1126/science.1262110 (2015).

42 Attig, J. et al. Splicing repression allows the gradual emergence of new Alu-exons in primate evolution. eLife 5, doi:10.7554/eLife.19545 (2016).

43 Wagner, A. Robustness and Evolvability in Living Systems. (Princeton University Press, 2007).

44 Tyner, C. et al. The UCSC Genome Browser database: 2017 update. Nucleic acids research 45, D626–D634, doi:10.1093/nar/gkw1134 (2017).

45 Li, W. & Freudenberg, J. Characterizing regions in the human genome unmappable by next-generation-sequencing at the read length of 1000 bases. Computational biology and chemistry 53 Pt A, 108–117, doi:10.1016/j.compbiolchem.2014.08.015 (2014).

46 Pickrell, J. K., Gaffney, D. J., Gilad, Y. & Pritchard, J. K. False positive peaks in ChIP-seq and other sequencing-based functional assays caused by unannotated high copy number regions. Bioinformatics 27, 2144–2146, doi:10.1093/bioinformatics/btr354 (2011).

47 Zhao, X., Valen, E., Parker, B. J. & Sandelin, A. Systematic clustering of transcription start site landscapes. PloS one 6, e23409, doi:10.1371/journal.pone.0023409 (2011).

48 Cohen, N. M., Kenigsberg, E. & Tanay, A. Primate CpG islands are maintained by heterogeneous evolutionary regimes involving minimal selection. Cell 145, 773–786, doi:10.1016/j.cell.2011.04.024 (2011).

49 Loytynoja, A. & Goldman, N. Phylogeny-aware gap placement prevents errors in sequence alignment and evolutionary analysis. Science 320, 1632–1635, doi:10.1126/science.1158395 (2008).

50 Ashkenazy, H. et al. FastML: a web server for probabilistic reconstruction of ancestral sequences. Nucleic acids research 40, W580–584, doi:10.1093/nar/gks498 (2012).

51 Guo, F. et al. The Transcriptome and DNA Methylome Landscapes of Human Primordial Germ Cells. Cell 161, 1437–1452, doi:10.1016/j.cell.2015.05.015 (2015).

52 International HapMap Consortium et al. A second generation human haplotype map of over 3.1 million SNPs. Nature 449, 851–861, doi:10.1038/nature06258 (2007).

53 Pereira, V. Automated paleontology of repetitive DNA with REANNOTATE. BMC genomics 9, 614, doi:10.1186/1471-2164-9-614 (2008).

54 Stempor, P. & Ahringer, J. SeqPlots - Interactive software for exploratory data analyses, pattern discovery and visualization in genomics. Wellcome open research 1, 14, doi:10.12688/wellcomeopenres.10004.1 (2016).

55 Ramirez, F., Dundar, F., Diehl, S., Gruning, B. A. & Manke, T. deepTools: a flexible platform for exploring deep-sequencing data. Nucleic acids research 42, W187–191, doi:10.1093/nar/gku365 (2014).

56 Grubert, F. et al. Genetic Control of Chromatin States in Humans Involves Local and Distal Chromosomal Interactions. Cell 162, 1051–1065, doi:10.1016/j.cell.2015.07.048 (2015).

57 Maurano, M. T. et al. Large-scale identification of sequence variants influencing human transcription factor occupancy in vivo. Nature genetics 47, 1393–1401, doi:10.1038/ng.3432 (2015).

58 Schultz, M. D. et al. Human body epigenome maps reveal noncanonical DNA methylation variation. Nature 523, 212–216, doi:10.1038/nature14465 (2015).

59 Tehranchi, A. K. et al. Pooled ChIP-Seq Links Variation in Transcription Factor Binding to Complex Disease Risk. Cell 165, 730–741, doi:10.1016/j.cell.2016.03.041 (2016).

